# Cytidine triphosphate promotes efficient ParB-dependent DNA condensation by facilitating one-dimensional spreading from *parS*

**DOI:** 10.1101/2021.02.11.430778

**Authors:** Francisco de Asis Balaguer, Clara Aicart-Ramos, Gemma LM Fisher, Sara de Bragança, Cesar L. Pastrana, Mark S. Dillingham, Fernando Moreno-Herrero

## Abstract

Faithful segregation of bacterial chromosomes relies on the ParABS partitioning system and the SMC complex. In this work, we used single molecule techniques to investigate the role of cytidine triphosphate (CTP) binding and hydrolysis in the critical interaction between centromere-like *parS* DNA sequences and the ParB CTPase. Using a combined dual optical tweezers confocal microscope, we observe the specific interaction of ParB with *parS* directly. Binding around *parS* is enhanced 4-fold by the presence of CTP or the non-hydrolysable analogue CTPγS. However, ParB proteins are also detected at a lower density in distal non-specific regions of DNA. This requires the presence of a *parS* loading site and is prevented by roadblocks on DNA, consistent with one dimensional diffusion by a sliding clamp. Magnetic tweezers experiments show that the spreading activity, which has an absolute requirement for CTP binding but not hydrolysis, results in the condensation of *parS*-containing DNA molecules at low nanomolar protein concentrations. We propose a model in which ParB-CTP-Mg^2+^ complexes move along DNA following loading at *parS* sites and protein:protein interactions result in the localised condensation of DNA within ParB networks.

## INTRODUCTION

In bacterial cells, the separation of sister chromosomes is performed by the ParABS system and the SMC complex (Marbouty et al., 2015; Song and Loparo, 2015; Wang et al., 2014). The ParABS system consists of the ATPase protein ParA, the DNA-binding protein ParB, and a centromere-like palindromic DNA sequence named *parS* (Funnell, 2016; Lin and Grossman, 1998). *In vivo* imaging experiments in *Bacillus subtilis* showed that multiple ParB proteins co-localize with SMC complexes at a given *parS* site forming distinctive clusters in the cell (Gruber and Errington, 2009; Sullivan et al., 2009). Notably, chromatin immuno-precipitation (ChIP) experiments indicate that ParB covers regions of up to 18 kilobase pairs (kbp) of DNA surrounding *parS* (Breier and Grossman, 2007; Graham et al., 2014; Minnen et al., 2016; Murray et al., 2006; Rodionov et al., 1999), a phenomenon named *spreading*. Originally, this spreading was interpreted as the formation of a nucleoprotein filament extending from a *parS* nucleation site (Murray et al., 2006; Rodionov et al., 1999). However, it later became clear that ParB foci contained far too few proteins to coat tens of kbp-long DNA segments (Graham et al., 2014). Instead, we and others have shown that ParB can self-associate to form networks which include specific binding to *parS* sequences but also non-specific binding to distal DNA segments. Overall, this results in the condensation and bridging of DNA at low forces (below 1 pN) (Fisher et al., 2017; Graham et al., 2014; Madariaga-Marcos et al., 2019, 2018; Taylor et al., 2015), and could explain how distant regions of DNA are bound by limited numbers of ParB proteins as shown in ChIP experiments. However, DNA condensation has only been observed *in vitro* at high ParB concentrations, in the low micromolar range. Moreover, and unexpectedly, *parS* sequences did not affect DNA condensation under these conditions (Taylor et al., 2015). Therefore the mechanism of ParB spreading and condensation, and in particular the molecular basis for the specific localisation around *parS*, has remained unclear despite extensive investigation *in vivo, in vitro* and *in silico* (Broedersz et al., 2014; Guilhas et al., 2020; Sanchez et al., 2015; Walter et al., 2020).

ParB proteins comprise three distinct domains (**Figure 1A and 1B**). The N-terminal domain (NTD) binds ParA (Bouet and Funnell, 1999; Davis et al., 1992; Radnedge et al., 1998; Vecchiarelli et al., 2010), and was recently appreciated to contain a CTP binding pocket that serves as a CTP-dependent dimerization interface (Osorio-Valeriano et al., 2019; Soh et al., 2019). Mutation R80A in the CTP binding pocket has been shown to impair nucleoid segregation and ParB spreading (Autret et al., 2001; Breier and Grossman, 2007). A central DNA binding domain (CDBD) binds specifically to the palindromic *parS* sequence and may also facilitate dimerization (Leonard et al., 2004; Schumacher and Funnell, 2005). In this central region, the mutation R149G within a helix-turn-helix motif impedes *parS* binding (Autret et al., 2001; Fisher et al., 2017; Gruber and Errington, 2009). Finally, the C-terminal domain (CTD) forms dimers with a lysine-rich surface that binds to DNA non-specifically (Fisher et al., 2017). Interactions between CTDs are essential for condensation *in vitro* and for the formation of ParB foci *in vivo* (Fisher et al., 2017). Based upon these results we proposed a model for DNA condensation dependent on ParB CTD dimerization and non-specific DNA (nsDNA) binding (Fisher et al., 2017). Further work using lateral pulling of long DNA molecules in a magnetic tweezers (MT) setup combined with total internal reflection fluorescence microscopy (TIRFM) demonstrated that ParB networks are highly dynamic and display a continual exchange of protein-protein and protein-DNA interfaces (Madariaga-Marcos et al., 2019).

**Figure 1.**
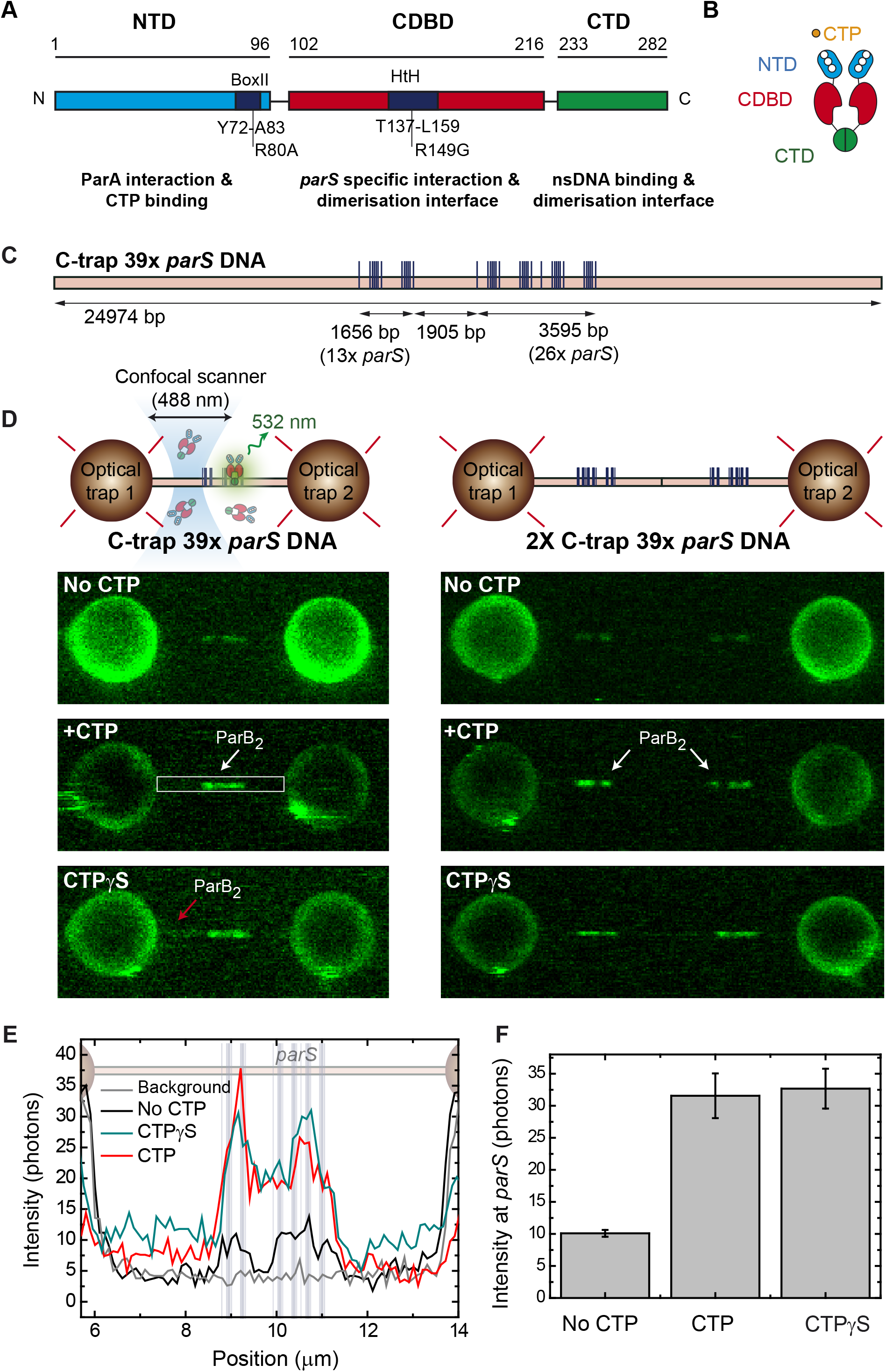
Direct visualisation of ParB specific binding to parS sites. (**A**) Domains and functional motifs of ParB as reported previously (Bartosik et al., 2004; Kusiak et al., 2011). Mutations R80A, defective for CTP binding, and R149G, defective for *parS* binding, are indicated. (**B**) ParB dimer cartoon showing dimerization through the central and C-terminal domains. The nucleotide binding site at the NTD is also indicated. (**C**) Schematic representation of the single-length 39x *parS* DNA used for C-trap experiments. The DNA contains 39 *parS* sequences distributed in 6 groups forming two clusters separated by 1905 bp (39x *parS* DNA). The positions of the *parS* sites in the DNA cartoon are represented to scale. (**D**) Schematic of the C-trap experiment where single and tandem (double-length) tethers are immobilised between two beads and scanned with a confocal microscope using 488 nm illumination (upper part). Representative confocal images of the experiment under no CTP, 2 mM CTP, or 2 mM CTPγS conditions (lower part) and 20 nM ParB_2_^AF488^. Dark to bright regions correspond to a scale of 0-30 photon counts for single-length tethers and 0-50 counts for tandem tethers. (**E**) Representative profiles (500 nm width) of the fluorescence intensity along the DNA axis of the confocal images depicted in **D** (only single-length tether data). Positions of the *parS* sequences are included to scale in the background. Brighter regions between the beads correlate with the position of the *parS* clusters. ParB proteins are also observed outside the *parS* region (red arrow) and in general the fluorescence intensity outside the *parS* region is always above the background and larger in CTPγS compared to CTP experiments. (**F**) Quantification of fluorescence intensity at the *parS*-containing region under no CTP, CTP and CTPγS conditions.

Recently, two laboratories demonstrated independent that *B. subtilis* ParB and the *Myxococcus xanthus* ParB hydrolyse cytidine triphosphate (CTP) to cytidine diphosphate (CDP) (Osorio-Valeriano et al., 2019; Soh et al., 2019) and require CTP for partition complex formation *in vivo* (Osorio-Valeriano et al., 2019). Importantly, *parS* DNA stimulates the binding and hydrolysis of CTP (Osorio-Valeriano et al., 2019; Soh et al., 2019), CTP has been shown to be necessary for ParB spreading *in vitro* (Jalal et al., 2020) and a model has been proposed in which centromeres assemble via the loading of ParB-DNA sliding clamps at *parS* sequences (Jalal et al., 2020; Soh et al., 2019). These observations fundamentally change our understanding of how ParB can become engaged with non-specific DNA surrounding *parS*, but the significance of CTP-dependent spreading in the formation of the ParB networks that cause DNA bridging and condensation has not been addressed.

Here, we investigated the role of CTP binding and hydrolysis in the binding of ParB to *parS* sequences and non-specific DNA at the single-molecule level. We present the first visualisation of the specific binding of ParB to *parS*. ParB proteins were also detected at non-specific DNA far from *parS*, albeit at a much lower density. Importantly, this only occurred in *parS*-containing DNA, suggesting spreading from *parS* sites. The placement of tight-binding protein roadblocks on DNA constrained the spreading, suggesting it arises from one-dimensional movement along the contour of DNA, consistent with a sliding clamp model. CTP binding mediated by a Mg^2+^ cofactor, but not CTP hydrolysis, was critical for spreading from *parS* to non-*parS* sites which triggered DNA condensation at nanomolar protein concentration. We propose a model where ParB-CTP-Mg^2+^ loads to *parS*, diffuses to non-*parS* sites and then self-associates, resulting in the condensation of kilobase-pair long DNA molecules.

## RESULTS

### Direct visualisation of the specific binding of ParB to *parS* sequences

In our previous work, performed in the absence of CTP, we used a combination of TIRFM and DNA stretching (by either flow or magnetic tweezers) to visualise binding of ParB to long DNA molecules (Madariaga-Marcos et al., 2019). Although specific ParB-*parS* interactions have been detected in electrophoretic mobility shift assays (EMSA) (Taylor et al., 2015), they have not yet been observed directly. Instead, regardless of the presence or absence of *parS* sequences, we and others observe a uniform non-specific coating of DNA with ParB accompanied by rapid DNA bridging and condensation (Graham et al., 2014; Madariaga-Marcos et al., 2019).

In this study, we investigated the effect of CTP on the specific and non-specific binding of ParB to *parS* using a fluorophore-conjugated ParB^AF488^, which retains specific and non-specific DNA binding activity (Madariaga-Marcos et al., 2019), and an experimental setup that combines confocal fluorescence microscopy with dual optical tweezers (C-trap, Lumicks) (Candelli et al., 2011; Newton et al., 2019) (see Methods, **Figure 1 and Figure 1 –figure supplement 1**). In this approach, DNA molecules are perpendicular to the optical axis of the microscope providing a homogeneous illumination along the molecule and confocal imaging provides a very high signal-to-noise ratio (albeit with a limited spatial resolution of around 250 nm). Additionally, the force applied to DNA is better defined and more uniform than in flow-stretch experiments.

Single DNA molecules were immobilized between two polystyrene beads and extended to almost their contour length by a force of ∼20 pN. To amplify the potential signal from specific binding to *parS* sites, we used a DNA substrate that contains 39 copies of the partially degenerated *parS* sequence (5’-TGTTCCACGTGAAACA) (Breier and Grossman, 2007; Taylor et al., 2015) arranged in two clusters (**Figure 1C** and **Figure 1 – figure supplement 2** and **3**). Then, we incubated the DNA with 20 nM ParB^AF488^ and took confocal images of the region of interest, including the beads as a reference, in the presence and absence of CTP-Mg^2+^ (**Figure 1D**). The images clearly showed two bright regions, one larger than the other, corresponding to the two *parS* clusters separated by 1905 bp. Note that, due to the design of the molecule (see Methods), double-length substrates can also be generated and trapped between the beads. In this case, the number of bright clusters were doubled, as expected (**Figure 1D**). Fluorescence intensity profiles of ParB correlated with the position of the *parS* clusters (**Figure 1E**). Importantly, ParB binding to *parS* did not require CTP in agreement with previous EMSA assays (Taylor et al., 2015) and bio-layer interferometric analysis (Jalal et al., 2020). However, the presence of CTP enhanced the fluorescence intensity within the *parS* clusters by 3-4 fold compared to in the absence of CTP (**Figure 1F**). Previously, we showed that the ParB binding equilibrium is established rapidly (within tens of seconds, (Madariaga-Marcos et al., 2019)) compared to our incubation time before confocal imaging. Therefore, these images represent the steady-state occupancy of ParB on DNA and the higher fluorescence intensity measured in the CTP case reflects a greater number of ParB molecules bound at or around the *parS* sequences compared to the no CTP condition. It is formally possible that the higher fluorescence measured in the presence of CTP could reflect a fluorescence enhancement effect, but this is unlikely since the ParB labelling site (S68C) is on a surface-exposed loop that is distant from the buried CTP molecules (Soh et al., 2019). In some images, a faint fluorescence signal was also observed outside the *parS* region (see red arrow in **Figure 1D**) in the CTP and CTPγS conditions and we will return to this point later. Control experiments with non-*parS* DNA did not show any protein binding at this ParB concentration (see below). This is the first direct visualization of ParB association specifically and precisely at *parS* sequences, and shows that this interaction is mediated by CTP binding in agreement with recently published works (Jalal et al., 2020; Osorio-Valeriano et al., 2019; Soh et al., 2019).

Next, we investigated the dynamics of the ParB-DNA interaction by taking kymographs with CTP/CTPγS or in the absence of nucleotide (see methods). The fluorescence intensity at the *parS* sites decayed with time but ParB remained visible for 30 s in both CTP/CTPγS conditions (**Figure 1-figure supplement 4A and 4B**). This helped to reveal the positioning of ParB relative to the *parS* sites throughout the 30 s kymograph (**Figure 1-figure supplement 4C**). The 39x*parS* DNA substrate includes two sequence clusters separated by 1905 bp, the smaller of which contains two groups of closely-spaced *parS* sequences, and the larger of which contains four such groups (**Figure 1-figure supplement 4C**). The 30 s average intensity profile clearly distinguished 6 foci corresponding to groups of ParB molecules precisely at their expected positions. The gradual decay of the fluorescence over tens of seconds (**Figure 1-figure supplement 4D**) indicates that the photobleaching kinetics are faster than the rates of ParB binding and unbinding. If the opposite were true then efficient protein turnover would result in a constant fluorescence level as was observed in our previous experiments performed at much higher ParB concentration (Madariaga-Marcos et al., 2019). Interestingly, the fluorescence at the *parS* clusters decayed marginally more slowly in the presence of CTPγS compared to CTP (**Figure 1-figure supplement 4D**).

### ParB spreading from *parS* sites occurs by sliding and requires CTP binding but not hydrolysis

With the aim of exploring ParB spreading from *parS* sites to distal DNA sites, we performed experiments in which we incubated the ParB protein with the DNA for 2 minutes before illuminating the sample (**Figure 2A**). By doing this, we prevented photobleaching of the proteins in the process of loading and spreading. We then compared the fluorescence intensity profiles of the first images obtained after incubation under different experimental conditions (**Figure 2B)**. Importantly, ParB proteins were now more clearly identified outside of the *parS* region (compared to **Figure 1D**), but only under CTP or CTPγS conditions (**Figure 2B**). Indeed, ParB proteins were sparsely distributed along the entire non-specific region (i.e., they did not only accumulate at or near the *parS* cluster) (red arrow, **Figure 2B and 2C**). The intensity in this non-*parS* region decayed within 1-2 seconds; an apparently shorter timescale than in the *parS* regions (**Video 1**), re-enforcing the idea that protein turnover is slow compared to photobleaching. Crucially, a control experiment using a non-*parS* DNA showed no protein bound at all in the presence of CTP (**Figure 2B and 2C**), supporting the notion that proteins located outside *parS* reached that location through the *parS* entry site.

**Figure 2.**
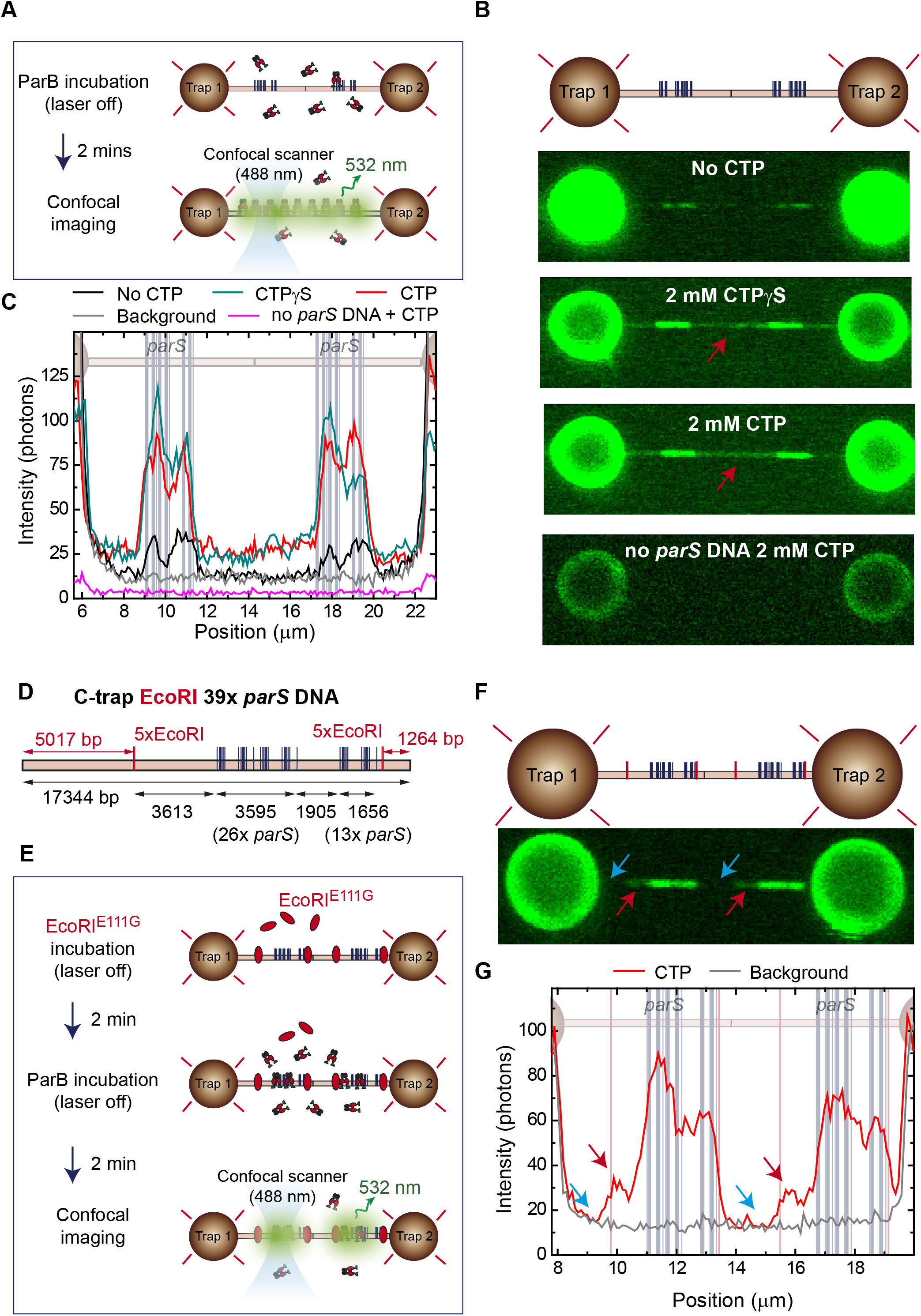
CTP binding promotes ParB spreading from parS. (**A**) Cartoon of the experiment. First, a tandem 39x *parS* DNA molecule is incubated with 20 nM ParB2 and 2 mM CTP-Mg^2+^. Then, following a 2 min incubation, the confocal laser is turned on and confocal images are taken. (**B**) Representative confocal images taken after 2 min ParB incubation in the dark using tandem 39x *parS* DNA under no CTP, CTP, or CTPγS conditions, as well as *parS*-free DNA (lambda DNA) and 2 mM CTP-Mg^2+^. ParB appears in non-*parS* regions only when using *parS* DNA and under CTP or CTPγS conditions (red arrows). Dark to bright regions correspond to a scale of 0-50 photon counts for *parS* DNA tethers and 0-25 counts for lambda DNA. (**C**) Corresponding average profiles (500 nm width) of the fluorescence intensity taken along the DNA axis of the confocal images. Positions of the *parS* sequences are included to scale in the background. (**D**) Schematic representation of the single-length EcoRI 39x *parS* DNA used for C-trap roadblock experiments. The DNA contains 39 *parS* sequences arranged as in **Figure 1C**, but also includes two groups of 5xEcoRI sites flanking the *parS* region. Note that one of the 5xEcoRI groups is located 3613 bp away from the last *parS* sequence, potentially allowing spreading from the *parS* region. The positions of the *parS* sites in the DNA cartoon are represented to scale. (**E**) Cartoon of the roadblock experiment designed to limit ParB spreading using the EcoRI^E111G^ mutant as a roadblock. The experiment is identical to that described in **A**, but first includes a 2 min pre-incubation with 100 nM EcoRI^E111G^, which is capable of DNA binding to EcoRI sites but unable to cleave the DNA, thus acting as a roadblock. (**F**) Confocal image showing limited spreading due to EcoRI^E111G^ blocking in tandem EcoRI 39x *parS* DNA. Brighter regions correspond to *parS* binding and the two dimmed regions correspond to limited spreading up to the EcoRI sites (red arrows). Regions inaccessible to ParB spreading are indicated with blue arrows. (**G**) Corresponding average profile (500 nm width) of the fluorescence intensity taken along the DNA axis of the confocal image. Positions of the *parS* sequences and EcoRI sites are included to scale in the background. Red arrows indicate the limited spreading of ParB up to EcoRI sites. Blue arrows indicate inaccessible regions to ParB.

Previous *in vitro* and *in vivo* experiments have shown that ParB spreading is hindered by DNA binding protein “roadblocks” engineered close to *parS* (Murray et al., 2006; Rodionov and Yarmolinsky, 2004; Soh et al., 2019). Therefore, to investigate the mechanism by which ParB spreads to non-specific sites we fabricated a 17-kb DNA molecule that contains the same 39x *parS* cluster flanked by two groups of 5xEcoRI sites (**Figure 2D**). These sequences would act as roadblocks after binding of EcoRI^E111G^, a catalytically inactive variant of the EcoRI restriction enzyme which has been used as a model protein roadblock (**Figure 2E**) (Brüning et al., 2018; King et al., 1989). Note that, if movement of ParB occurs along the contour of DNA (i.e., by sliding from *parS*), then we expect ∼5-kbp and ∼1-kbp DNA segments of the molecule to remain free of ParB proteins. Additionally, the molecule contains a ∼3.6-kbp non-*parS* area between the last *parS* sequence of the 39x *parS* cluster and one of the 5xEcoRI sites which we would expect to become populated with ParB via a sliding mechanism (**Figure 2D**). The imaging experiments were performed as described in **Figure 2A** but included an additional incubation step with 100 nM EcoRI^E111G^, prior to incubation with ParB^AF488^. The confocal laser was turned on after the incubations with EcoRI^E111G^ and ParB^AF488^, and confocal images were obtained on a tandem EcoRI 39x *parS* DNA (**Figure 2F**). As expected, brighter regions correlated very well with the *parS* clusters. Importantly, a faint region also appeared flanking the *parS* cluster and bordered by the EcoRI^E111G^ roadblocks, which appeared to have constrained ParB spreading (see red arrows, **Figure 2F**). Indeed, fluorescence profiles showed high intensity associated with the *parS* region, lower intensity signals produced by ParB spreading from *parS*, (see red arrows, **Figure 2F and Figure 2G**), and no signal associated with regions protected by EcoRI sites (see blue arrows, **Figure 2F and Figure 2G**). Subsequent confocal images reflected the photobleaching of this region in contrast with the brighter *parS* area, confirming the low exchange of proteins outside *parS* (**Video 2**). Altogether, these experiments show that CTP binding promotes movement of ParB over kilobase-pair distances away from *parS* sites to non-specific regions of DNA. The fact that this movement can be constrained by protein roadblocks suggests that it occurs by sliding from *parS*.

### CTP binding dramatically enhances the *parS* sequence-specificity of ParB-dependent DNA condensation

We have previously shown that ParB condenses DNA and that the CTD plays an important role in this function (Fisher et al., 2017; Taylor et al., 2015). However, condensation occurred at micromolar protein concentration and was not apparently specific to *parS*-containing DNA. Now, we aimed to revisit these experiments in the light of the discovery of CTP as an important mediator of ParB-DNA interactions (Osorio-Valeriano et al., 2019; Soh et al., 2019). Note that the optical trap experiments described above were necessarily performed at forces which are non-permissive for condensation, to keep the DNA extended for optimal fluorescence visualization. We therefore switched to a magnetic tweezers setup which is more appropriate for low-force experiments (Taylor et al., 2015) (**Figure 3A**). Single DNA molecules containing a set of 13x *parS* sequences were immobilized between a glass surface and super-paramagnetic beads (**Figure 3B**). A pair of magnets were then employed to stretch the DNA and apply forces in the 0.1-5 pN range. The DNA was incubated with ParB at the higher force level for two minutes, and then the force was lowered to 0.33 pN, which is permissive for DNA condensation. The extension of the tether was monitored in real time leading to condensation time-course plots.

**Figure 3.**
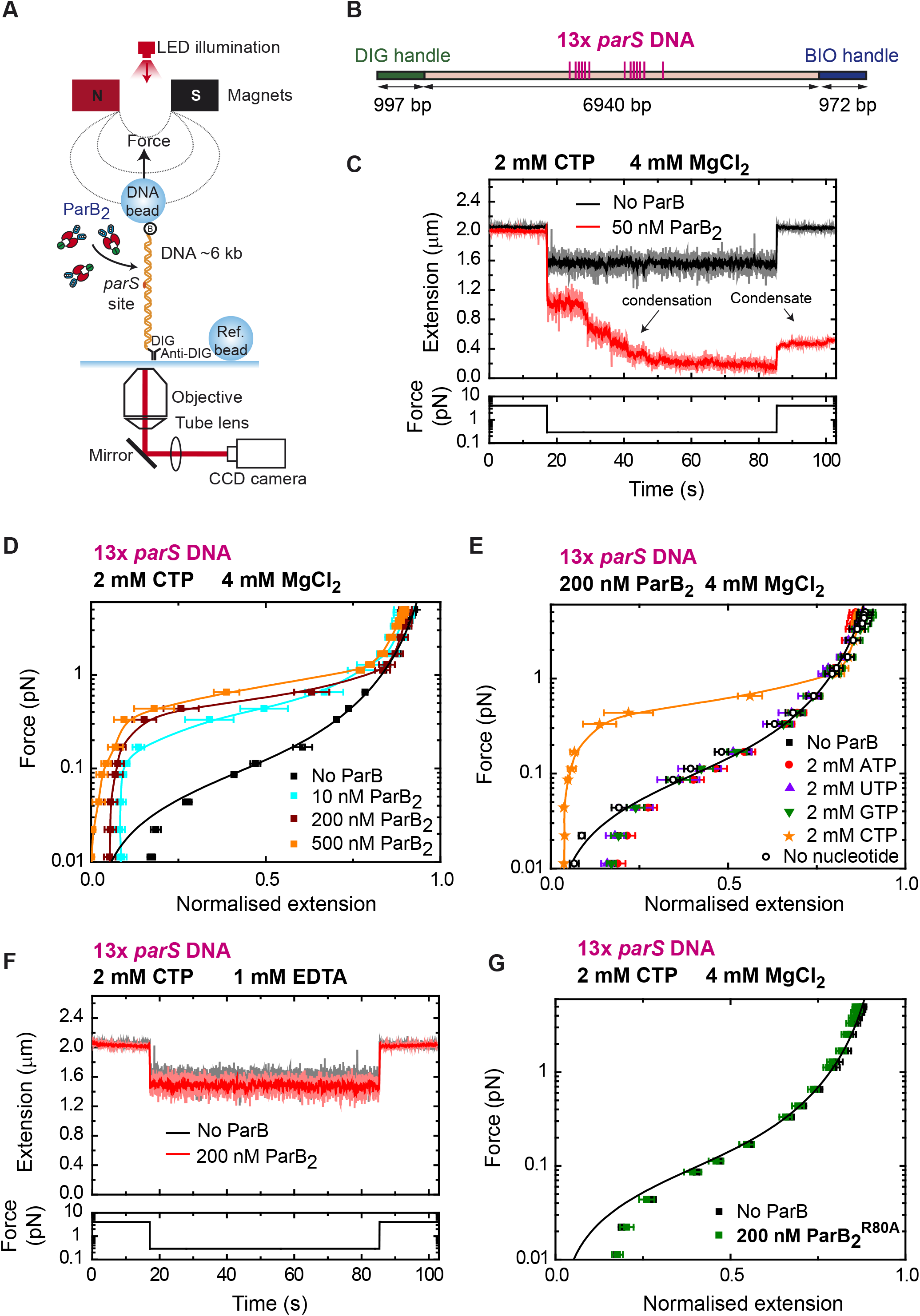
DNA condensation is induced by ParB at nanomolar concentrations in the presence of CTP. (**A**) Cartoon of the basic magnetic tweezers components and the layout of the experiment. (**B**) Schematic representation of the 13x *parS* DNA used for MT experiments. The positions of the *parS* sites in the DNA cartoon are represented to scale. (**C**) Condensation assay. DNA is held at 4 pN while 50 nM ParB2 is injected into the fluid cell in the presence of 2 mM CTP and 4 Mm MgCl_2_. Following a 2-min incubation, the force is lowered to 0.3 pN and the extension recorded (red data). The extension in the absence of protein is shown in black. DNA could not recover the original extension by force after condensation at low force. (**D**) Average force-extension curves of 13x *parS* DNA molecules in the presence of 2 mM CTP, 4 mM MgCl_2_ and increasing concentrations of ParB2. A concentration of only 10 nM ParB2 was able to condense the 13x *parS* DNA. (**E**) Average force-extension curves of 13x *parS* DNA taken under the stated conditions and in the presence of different nucleotides or with no nucleotide. Only CTP produces condensation of *parS* DNA. Solid lines in the condensed data are guides for the eye. Errors are standard error of the mean for measurements taken on different molecules (N ≥ 6). (**F**) Condensation assay of 13x *parS* DNA under 2 mM CTP and 1 mM EDTA conditions. DNA condensation by ParB and CTP requires Mg^2+^. (**G**) CTP-binding mutant, ParB^R80A^, does not condense 13x *parS* DNA under standard CTP-Mg^2+^ conditions. No ParB data represent force-extension curves of DNA taken in the absence of protein and are fitted to the worm-like chain model. Errors are standard error of the mean for measurements taken on different molecules (N = 7).

DNAs containing *parS* rapidly condensed in assays with 50 nM ParB2, CTP and Mg^2+^ (**Figure 3C**). In fact, DNA condensation was observed at even lower concentrations of 5-10 nM, but the rate was markedly slower (**Figure 3-figure supplement 1A**). In the presence of CTP-Mg^2+^, DNA was condensed by ParB at forces of up to 1 pN using only 10 nM protein (**Figure 3D**). This maximum condensation force was similar to that described in non-*parS* DNA using micromolar ParB concentrations (Taylor et al., 2015), suggesting a similar mechanism of condensation. Experiments using ATP, UTP or GTP did not produce any DNA condensation, confirming the specificity of ParB for CTP and linking CTP binding to condensation (**Figure 3E**).

The recent crystal structure of the *M. xanthus* ParB-like protein PadC showed that Mg^2+^ is a cofactor of CTP at the CTP binding site (Osorio-Valeriano et al., 2019). Therefore, we explored the role of Mg^2+^ in the ParB condensation function. ParB did not induce any condensation in a buffer containing 1 mM EDTA and 200 nM ParB2 (**Figure 3F**). Mutation of the R80 residue of ParB to alanine leads to loss of function in *B. subtilis* and impairs subcellular localization of ParB (Autret et al., 2001; Graham et al., 2014). Additionally, CTP binding is also abolished in *Bs*ParB^R80A^ (Soh et al., 2019). We therefore ask if this mutation might affect the condensation function of ParB. Indeed, magnetic tweezers experiments showed no DNA condensation by ParB^R80A^ under CTP-Mg^2+^ conditions (**Figure 3G**) supporting the idea that CTP binding is required for condensation at nanomolar ParB concentrations. In the absence of CTP, ParB was unable to condense *parS*-containing DNA unless its concentration was raised to the micromolar range, as reported previously (**Figure 3-figure supplements 1B and 1C**) (Taylor et al., 2015).

We next asked whether CTP hydrolysis is required for DNA condensation by exploiting the non-hydrolysable analogue CTPγS. We took time-courses in the presence of 2 mM CTPγS and obtained force-extension curves at different ParB concentrations (**Figure 4A**). We did not observe any significant difference in force-extension measurements compared to the CTP case (**Figure 3D**). The DNA was still condensed at nanomolar ParB and condensation was abolished by EDTA (**Figure 4B**). Altogether, we conclude that binding to both CTP and the Mg^2+^ cofactor are required for DNA condensation by nanomolar ParB, but that CTP hydrolysis is not.

**Figure 4.**
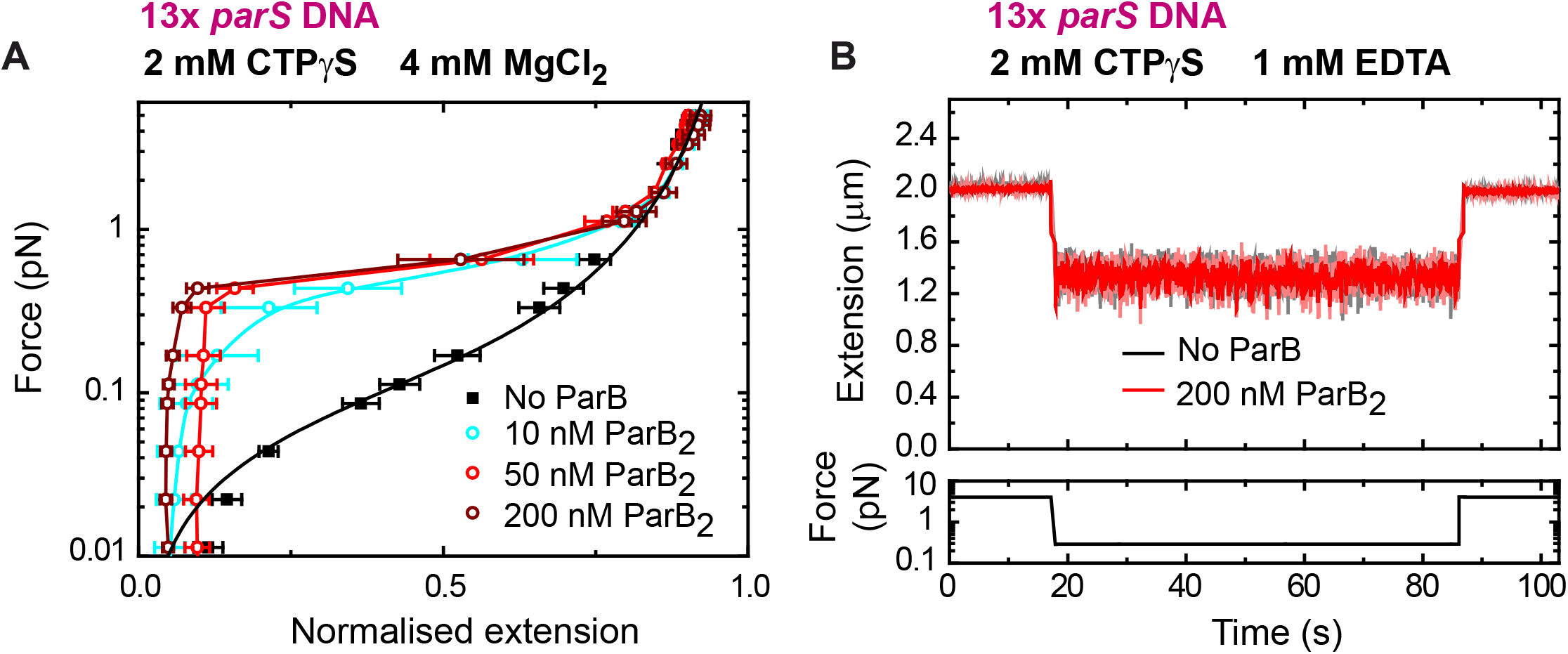
DNA condensation by nanomolar ParB requires CTP binding but not hydrolysis. (**A**) Average force-extension curves of 13x *parS* DNA molecules in the presence of 2 mM CTPγS, 4 mM MgCl_2_ and increasing concentrations of ParB. Results obtained with CTP (**Figure 3D**) and CTPγS were very similar. No ParB data represent force-extension curves of DNA taken in the absence of protein and are fitted to the worm-like chain model. Solid lines in condensed data are guides for the eye. Errors are standard error of the mean for measurements taken on different molecules (N ≥ 7). (**B**) Condensation assay of 13x *parS* DNA under 2 mM CTPγS and 1 mM EDTA conditions. DNA condensation by ParB and CTPγS requires Mg^2+^.

Finally, we investigated whether the dramatic stimulation of DNA condensation afforded by CTP was specific to *parS*-containing DNA and explored how condensation was affected by the number of *parS* sequences present in the substrate. Experiments using a DNA substrate with scrambled *parS* sequences did not show condensation even at ParB2 concentrations of 200 nM (**Figure 5A**). Moreover, to confirm that the DNA condensation was mediated by *parS* binding we used the mutant ParB^R149G^ that cannot bind *parS* (Autret et al., 2001; Fisher et al., 2017; Gruber and Errington, 2009). As expected, no condensation was observed using ParB^R149G^ under conditions proficient for condensation with wild type ParB (**Figure 5B**). We next obtained force-extension curves using substrates containing different numbers of *parS* sequences. DNA molecules containing from 1 to 26 *parS* sequences and similar overall length were fabricated (**Figure 5C and Figure 5E**, see methods). DNA condensation was observed in substrates with 26, 13, and 7 *parS* sequences with a clear correlation between the number of *parS* and the maximum force permissive for condensation (**Figure 5D**). Experiments with substrates containing 1, 2, or 4 copies of *parS* did not result in condensation even at a relatively high ParB2 concentration of 200 nM (**Figure 5E** and **Figure 5F**). These experiments indicated a requirement for a minimal number of *parS* sites (between 5 and 7) for condensation under these conditions.

**Figure 5.**
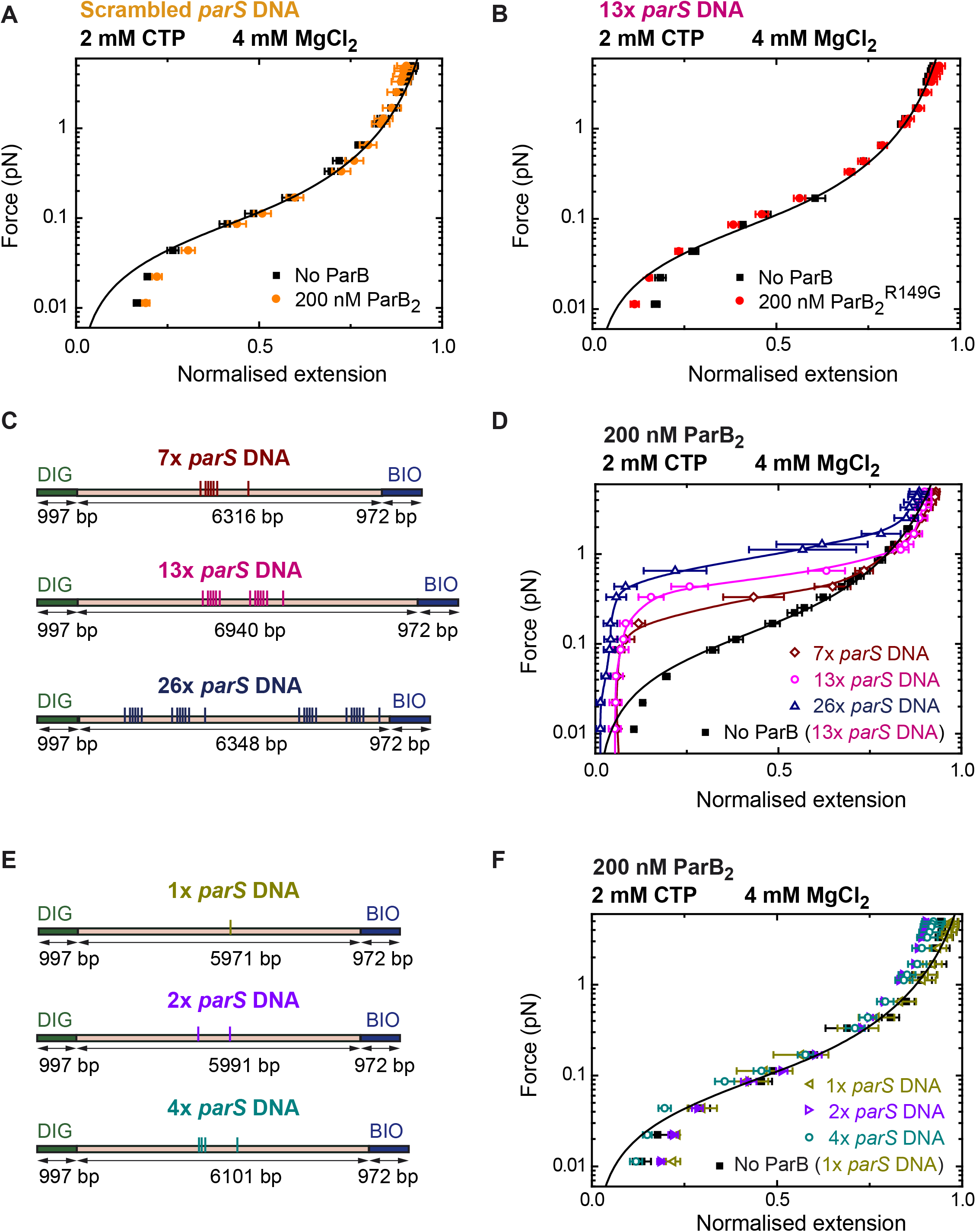
DNA condensation by nanomolar ParB is parS dependent. (**A**) ParB does not condense scrambled *parS* DNA under standard CTP-Mg^2+^ conditions. Errors are standard error of the mean of measurements on different molecules (N = 5). (**B**) The ParS-binding mutant, ParB^R149G^, does not condense 13x *parS* DNA under standard CTP-Mg^2+^ conditions. Errors are standard error of the mean of measurements on different molecules (N = 14). (**C**) Schematic representation of DNA substrates containing 7, 13, and 26 copies of *parS*. The positions of the *parS* sites in the DNA cartoon are represented to scale. (**D**) Average force-extension curves of 7x *parS* DNA, 13x *parS* DNA, and 26x *parS* DNA obtained under standard CTP-Mg^2+^ conditions. The condensation force correlates with increasing number of *parS* sequences. Solid lines in condensed data are guides for the eye. Errors are standard error of the mean of measurements on different molecules (N ≥ 7). (**E**) Schematic representation of DNA substrates containing 1, 2, and 4 copies of *parS*. The positions of the *parS* sites in the DNA cartoon are represented to scale. (**F**) Average force-extension curves of 1x *parS* DNA, 2x *parS* DNA, and 4x *parS* DNA obtained under standard CTP-Mg^2+^ conditions. No condensation was observed for these three experiments. Errors are standard error of the mean of measurements on different molecules (N ≥ 7). No ParB data represent force-extension curves of DNA taken in the absence of protein and are fitted to the worm-like chain model.

Together, these data show that CTP binding dramatically enhances DNA condensation by ParB such that it occurs efficiently at low nanomolar ParB concentration. Importantly, this stimulatory effect is completely specific for *parS*-containing DNA molecules as CTP does not improve the condensation of *parS*-free molecules that can be observed at high concentration of ParB. This presumably reflects the CTP- and *parS*-dependent recruitment of ParB sliding clamps that subsequently multimerise to effect bridging interactions between distal DNA segments.

## DISCUSSION

We report here the first visualisation of ParB binding to *parS* sequences. Previous observation of the specific binding to *parS* was hindered by the fact that ParB binding to DNA induces condensation and bridging (Graham et al., 2014; Taylor et al., 2015). To overcome this issue, we and others stretched the DNA molecules using flow or magnetic pulling combined with TIRF microscopy (Graham et al., 2014; Madariaga-Marcos et al., 2019). However, these experiments were performed in the absence of CTP and required high concentrations of ParB, conditions which allow ParB to interact directly (i.e., from free solution rather than via *parS* loading sites) with non-specific DNA. The recent discovery that ParB is a *parS*-dependent CTPase has led to the proposal of radically new models for ParB-DNA interactions in which *parS* acts as a loading site for ParB-DNA sliding clamps (Osorio-Valeriano et al., 2019; Soh et al., 2019). Together with bulk bio-layer interferometric analysis (Jalal et al., 2020), this work suggests that ParB binds to *parS* in the apo state and CTP-binding induces a conformational change that liberates the ParB dimer from *parS* allowing spreading. Motivated by these important new observations, we have revisited our earlier single molecule experiments using much lower concentrations of ParB in the presence of CTP. Our results confirm many aspects of the published models but also extend them, by addressing how CTP facilitates localised DNA condensation around *parS* sites.

In an attempt to visualise the specific binding of ParB to *parS* in the presence of CTP, Soh et al. observed stretched DNA bound to a glass surface using TIRF microscopy and reported accumulation of ParB around *parS* (Soh et al., 2019). Here, we used a combination of optical tweezers and confocal microscopy that allowed us not only to observe more precisely the direct binding of ParB to single DNA molecules, but also facilitated the quantification of fluorescence intensity around *parS* sites and distal non-specific sites. In our assay we found very good correlation between the position of ParB in time-averaged intensity profiles and the position of the groups of *parS* sites engineered into our substrate. These profiles easily allowed us to distinguish between the different orientations of substrate molecules (**Figure 1**). In these experiments, confocal illumination was initiated after a long ParB incubation with the substrate DNA such that the initial images should reflect equilibrium binding conditions (Madariaga-Marcos et al., 2019). In the absence of CTP, we observed a clear fluorescence intensity at *parS* sequences, and a zero intensity (within error) in distal regions of non-specific DNA. We interpret this as reflecting the highly specific binding of ParB in an open clamp conformation directly to the *parS* sequences via the HtH motifs in the CDBD domain (**Figure 6, step 1**).

**Figure 6.**
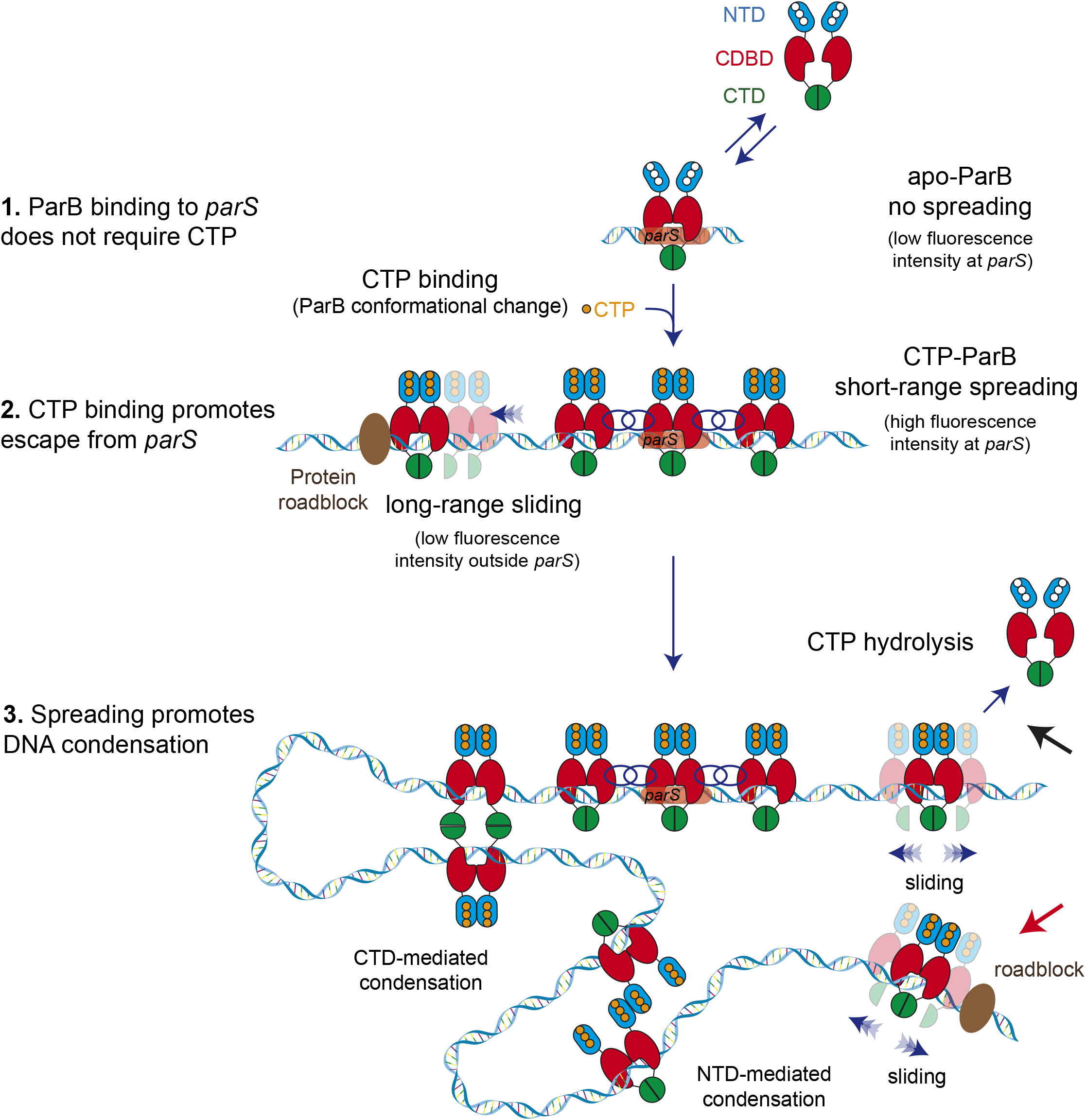
Model for ParB-dependent DNA condensation around parS sequences. (**Step 1**) ParB binding to *parS* does not require CTP as observed from C-trap experiments. *ParS*-bound apo-ParB does not spread from *parS*. (**Step 2**) CTP binding to ParB induces a conformational change to a sliding clamp which then escapes from *parS* to neighbouring non-specific DNA. Potential interactions between the ParB proteins around *parS* are represented by interlaced blue circles. Some ParB proteins are able to slide/diffuse long distances. (**Step 3**) ParB spreading and diffusion promotes the interaction with other CTP-ParB dimers through the CTD of ParB (Fisher et al., 2017), resulting in DNA condensation by forming large DNA loops. Alternatively, other protein-protein interaction such those mediated by the NTD (shown in figure) or the CDBD of ParB could result in DNA condensation. CTP hydrolysis might be a means to recover ParB dimers from the DNA (black arrow). Protein roadblocks constrain diffusion of ParB proteins (red arrow).

When we next added either CTP or the non-hydrolysable analogue CTPγS, we saw a 4-fold increased fluorescence intensity around the *parS* sequences and a lower (but clearly non-zero) intensity in distal non-specific regions of the DNA. Note that, because this is a single molecule experiment, the increased intensity associated with the *parS* clusters strongly suggests that there is a higher density of ParB dimers on the DNA at or near (i.e., within the spatial resolution of the imaging; ∼250 nm) *parS* sequences. We interpret this as the CTP binding-dependent conversion of *parS*-bound ParB dimers into sliding clamps that move into neighbouring regions of the DNA to allow additional ParB open clamps to be recruited to the DNA (**Figure 6, step 2**). It is interesting to note that the presence of CTP and CTPγS does not prevent the initial binding of ParB to DNA by favouring a closed clamp structure *before* DNA association. This might suggest either that the DNA entry gate for ParB is a different interface (e.g., transient opening of the CTD) or simply that the CTP cannot bind efficiently until after DNA enters ParB though the NTD entry gate. In either case, following the formation of closed ParB clamps on DNA, we imagine that they spread but remain largely restrained to the region of non-specific DNA immediately surrounding *parS* sequences as a result of protein:protein interactions (**Figure 6, step 2, interlaced circles**). In our C-trap experiment, the ends of the DNA substrate are held at the beads, meaning that the DNA cannot begin to condense by sliding into the ParB networks that are forming around *parS*. We will return to this point in our interpretation of the magnetic tweezers experiments below.

The weaker fluorescence intensity in distal non-specific regions of the substrate is observed exclusively in CTP or CTPγS conditions and is interpreted as ParB sliding clamps which have escaped from the ParB network at *parS* (**Figure 6, step 2**). Evidence that these ParB molecules are indeed involved in one-dimensional sliding interactions with the DNA is provided by roadblock experiments with a catalytically-inactive EcoRI mutant. The weaker fluorescence intensity associated with sparsely distributed ParB molecules in non-specific DNA regions was only observed in segments of DNA containing *parS* sequences and was corralled by EcoRI roadblocks placed either side of the centromere sequences (**Figure 6, step 2**).

Despite the fact that free ParB is always present in the imaging channel in our experiments, we observed a rapid decay of the fluorescence intensity upon illumination. This suggests that the photobleaching rate is faster than the binding and unbinding kinetics of ParB. Interestingly, we observed an apparently faster fluorescence loss at distal regions of non-specific DNA when compared to the regions close to *parS* (**Video 1 and 2**). This may reflect the much higher density of ParB that our model anticipates in *parS* regions compared to distal non-specific DNA. Alternatively, continuous loading at *parS* may counteract the fluorescence decay caused by photobleaching to a greater extent than at distal DNA regions (where ParB cannot bind from free solution). We also found that the fluorescence decay was marginally slower in experiments performed with CTPγS compared to CTP. This is consistent with the idea that the role of CTP hydrolysis is to open the ParB clamp to allow release from DNA and recycling.

In order to investigate the effects of CTP on ParB-induced DNA condensation we employed a magnetic tweezers apparatus. In the presence of CTP-Mg^2+^, ParB condensed *parS*-containing DNA at nanomolar concentrations, much lower than those reported before (Fisher et al., 2017; Taylor et al., 2015) (**Figure 3**). Importantly, this strong stimulatory effect of CTP on DNA condensation, which is completely specific for *parS*-containing DNA, helps to resolve the question of how ParB networks can form uniquely at *parS* loci within the huge bacterial chromosome. This had not been readily apparent from previous experiments performed without CTP (see (Graham et al., 2014; Madariaga-Marcos et al., 2019; Taylor et al., 2015) for discussion). Control experiments showed that efficient condensation required both the *parS* sequence and either CTP or CTPγS, suggesting that it is CTP binding but not its hydrolysis that is important for promoting condensation. Interestingly, we also found a *parS* sequence dose-dependence for condensation. The condensation force increased with the number of *parS* sites (**Figure 5**) and a minimum value of between 5 and 7 sites was required to observe condensation under the conditions used here.

We can interpret the behaviour observed in the magnetic tweezers using a simple extension of the model described above for the confocal microscopy experiments. In the magnetic tweezers experiment, the DNA ends are held apart by a very low force and the DNA is able to condense. Therefore, the ParB dimers which load around the *parS* sequence (**Figure 6, step 2**), or those which escape from the ParB network, can self-associate and draw DNA into the complex (**Figure 6, step 3**). This creates a dynamic network of ParB molecules constraining DNA loops. The greater the number of *parS* sequences present in the DNA, the greater will be the loading rate of ParB clamps into non-specific regions of the DNA, and the more stable will be the condensation against weak restraining forces. *In vivo* experiments have shown that a single *parS* is enough to promote chromosome segregation (Broedersz et al., 2014; Jecz et al., 2015; Wang et al., 2017). This might simply reflect the difference in restraining forces, ParB concentrations, or solution conditions that are found *in vivo* compared to our magnetic tweezers experiments. Alternatively, a failure to condense DNA over large distance scales does not necessarily imply the lack of formation of a ParB-*parS* complex which is sufficient for chromosome segregation.

The molecular basis for the protein-protein interactions that hold the ParB network together in our model remain unclear and are an important subject for future study. ParB contains multiple non-exclusive interfaces within all three domains that could be relevant to this activity (Song et al., 2017). For example, we have previously provided direct evidence that disruption of CTD-CTD interactions decondenses DNA *in vitro* and prevents the formation of ParB networks *in vivo*, so these protein-protein interactions may contribute to network formation (**Figure 6, step 3**). However, we note that our results can also be re-interpreted in the light of the new concept of topological engagement between DNA and a ParB toroidal clamp. If CTD-CTD interactions are important for closing the sliding clamp around DNA as has been suggested (Jalal et al., 2020; Soh et al., 2019), then disruption of these interactions would dissolve ParB networks, not by breaking protein-protein interactions between ParB molecules, but rather by releasing the constrained DNA loops and promoting ParB dissociation into free solution. Other possible interfaces that may establish ParB networks include those that have been observed between the CDBD or NTD (Chen et al., 2015; Leonard et al., 2004; Schumacher and Funnell, 2005). A final open question concerns the loading of the SMC complexes at bacterial centromeres which leads to the individualisation of bacterial chromosomes following DNA replication (Hayes and Barillà, 2006; Schumacher, 2008). Despite the intimate functional relationship between the ParABS partitioning system and SMC complexes, the nature of the physical interactions between these systems and their regulation are not understood.

## MATERIALS AND METHODS

### Protein preparation

WT-ParB, R149G ParB and AF488-ParB were prepared as described (Fisher et al., 2017; Madariaga-Marcos et al., 2019). An expression construct for R80A ParB was generated by site-directed mutagenesis of the wild type expression plasmid (QuikChangeII XL, Agilent Technologies). The mutant protein was expressed and purified in the same manner as wild type. The EcoRI E111G variant was a gift from Michelle Hawkins (University of York) and was prepared as described previously (King et al., 1989).

### Fabrication of DNA plasmids with multiple copies of *parS*

DNA plasmids containing multiples *parS* sequences (optimal sequence of *B. subtilis parS* = 5’-TGTTCCACGTGAAACA) were produced by modification of the plasmids described in (Taylor et al., 2015), where the cloning of a plasmid containing a ‘scrambled’ *parS* site (scrambled *parS*: 5’-**C**GT**G**CC**CA**G**G**GA**G**ACA; bold represents mutated nucleotides) was also reported.

Plasmids with increasing number of *parS* sequences were produced as follows. First, we annealed two long oligonucleotides (**Table S1**) containing 2 *parS* sites separated by a single XbaI restriction site. The oligonucleotides were hybridized by heating at 95°C for 5 min, and cooled down to 20°C at a -1°C min^-1^ rate in hybridization buffer (10 mM Tris-HCl pH 8.0, 1 mM EDTA, 200 mM NaCl and 5 mM MgCl_2_). These oligonucleotides were designed to create an incomplete XbaI site at both ends after ligation, so that once ligated to a cloning plasmid they cannot be cut again by XbaI. The single bona-fide XbaI site located in the middle of the oligonucleotide insert allows repetition of the ligation process in the cloning plasmid as many times as desired to add new pairs of *parS* sequences. Plasmids containing 1x *parS*, 2x *parS*, 4x *parS*, and 7x *parS* were obtained following this procedure. Inserts for subsequent cloning of plasmids containing 13x *parS*, 26x *parS*, and 39x *parS* were produced by PCR using a high-fidelity polymerase (Phusion Polymerase, NEB) (see **Table S1** for primer sequences). Plasmids were cloned in DH5α Competent cells (ThermoFisher Scientific) and purified from cultures using a QIAprep Spin Miniprep Kit (QIAGEN). All plasmids were checked by DNA sequence analysis. A detailed description of these procedures is included in **Supplementary Methods** and in **Figure 1 – figure supplement 2**.

### Fabrication of large plasmids (> 17 kbp) for C-trap experiments

Long fragments of DNA (> 17 kbp) containing a custom sequence for C-trap experiments were fabricated as follows. A large plasmid was formed by ligation of three pieces. Two of them correspond to two PCR-fabricated DNAs (C1 and C2) derived from lambda DNA (NEB) as template and including suitable restriction sites in the designed primers (**Table S1**). The third part (C3) containing the sequence of interest (e.g., multiple copies of the *parS* sequence) was produced either by PCR or plasmid digestion. The three fragments were then ligated and DH5α competent cells transformed by regular heat shock procedure. Large plasmids were purified from cultures using QIAprep Spin Miniprep Kit and checked by DNA sequence analysis. A detailed description of these procedures is included in **Supplementary Methods** and in **Figure 1 – figure supplement 3**.

Plasmids containing the *parS* region flanked by two clusters of 5x EcoRI restriction sites were produced following the same procedure but replacing parts C1 and C2 with fragments C1-EcoRI and C2-EcoRI, each one including a cluster of 5x EcoRI restriction sites at the desired position (**Table S1**).

### Magnetic tweezers DNA substrates

MT DNA substrates were produced as described in (Taylor et al., 2015) and essentially consist of a central part (∼6 kbp) containing different number of *parS* sequences or a non-specific scrambled *parS* site, flanked by two smaller fragments (∼1 kbp) labelled with biotins or digoxigenins used as immobilization handles. Handles for MT substrates were prepared by PCR (see **Table S1** for primers) including 200 μM final concentration of each dNTP (G,C,T,A) and 10 µM Bio-16-dUTP or Dig-11-dUTP (all from Roche). Labelled handles specifically bind either to a glass surface covered with anti-digoxigenins or to superparamagnetic beads covered with streptavidin. About 40% of molecules fabricated using this procedure were torsionally constrained in MT experiments. Sequences of the central part of magnetic tweezers substrate are included in **Supplementary Methods**.

### C-trap DNA substrates

C-trap DNA substrates consisted of a large central part of ∼17 kbp (EcoRI 39x *parS* DNA) or ∼25 kbp (39x *parS* DNA) containing 39 copies of the *parS* sequence, flanked or not by 2 clusters of 5x EcoRI restriction sites, respectively, were produced by linearization of large plasmids with NotI (NEB). Without further purification, the fragment was ligated to highly biotinylated handles of ∼1 kbp ending in *NotI*. Handles for C-trap substrates were prepared by PCR (see **Table S1** for primers) including 200 μM final concentration of each dNTP (G,C,A), 140 µM dTTP, and 66 µM Bio-16-dUTP. These handles were highly biotinylated to facilitate the capture of DNA molecules in C-trap experiments. As both sides of the DNA fragment end in *NotI*, it is possible to generate tandem (double-length) tethers of ∼34 kbp or ∼50 kbp (tandem EcoRI 39x *parS* DNA or 39x *parS* DNA, respectively) flanked by the labelled handles. Sequences of the central part of C-trap substrates are included in **Supplementary Methods**.

A control C-trap DNA substrate based on lambda DNA was prepared according to a previously described protocol (Wasserman et al., 2020) with slight modifications. 10 nM lambda DNA was incubated with 33 μM each of dGTP, dATP, biotin-16-dUTP and biotin-14-dCTP (Thermo Fisher), and 5 units of DNA polymerase (Klenow Fragment (3’→5’ exo-), NEB) in 1X NEB2 buffer for 1 hour at 37°C. Reaction was followed by heat inactivation of the enzyme for 20 min at 75°C. Sample was ready to use in C-trap experiments without further purification. DNAs were never exposed to intercalanting dyes or UV radiation during their production and were stored at 4°C.

### Measurement of NTP hydrolysis by Malachite Green colorimetric detection

A pair of oligonucleotides containing a *Scrambled parS* site, 1x *parS* or 2x *parS* sites (see **Table S1** for sequences) were hybridised by heating at 95°C for 5 min, and cooled down to 20°C at a - 1°C min^-1^ rate in hybridization buffer (10 mM Tris-HCl pH 8.0, 1 mM EDTA, 200 mM NaCl and 5 mM MgCl_2_). Mixtures of NTP (2x) and DNA (2x) in reaction buffer (100 mM NaCl, 50 mM Tris-HCl pH 7.5, 4 mM MgCl_2_, 1 mM DTT and 0.1% Tween-20) were prepared on ice. Protein solutions (2x) containing either wild type ParB or ParB^AF488^ in reaction buffer were also prepared on ice. NTP/DNA pre-mix (5 μl) was added to protein solution (5 μl) and mixed on ice. Phosphate standards and blanks were prepared in parallel for each experiment. After mixing, samples containing 1 mM NTP, 0.5 μM DNA, 8 μM ParB2 were placed in a PCR machine set to 25 °C for 30 min. Additionally, different concentrations (0.25 μM, 0.75 μM, and 1 μM) of a DNA with 2 *parS* sequences were tested in presence of 1 mM CTP. Samples were diluted by the addition of 70 μL water, then mixed with 20 μL working reagent (WR) (Sigma, Ref MAK 307) and transferred to a flat bottom 96 well plate. The plate was incubated for 30 minutes at 25 °C and the absorbance was measured at a wavelength of 620 nm in a SpectraMax® iD3 (VERTEX Technics) plate reader that uses the SoftMax® Pro7 software. Readings were performed in rounds to preserve the same 30 min WR incubation time for all samples. Absorbance values from the phosphate standard samples were corrected with the absorbance for 0 μM phosphate. Absorbance values from the ParB samples were corrected by the reaction buffer absorbance (blank). Absorbance values from the phosphate standard samples were used to plot an OD620 nm versus pmol phosphate standard curve. All samples were tested in duplicate. ParB^AF488^ retains *parS*-stimulated CTPase activity within 2-fold levels of wild type protein (**Figure 1 –figure supplement 1B and 1C**).

### Magnetic tweezers experiments

Magnetic tweezers assays were performed using a home-made setup similar to an apparatus that has been described previously (Carrasco et al., 2013; Kemmerich et al., 2016; Pastrana et al., 2016; Strick et al., 1998). Briefly, optical images of micrometer-sized superparamagnetic beads tethered to a glass surface by DNA constructs are acquired with a 100x oil-immersion objective and a CCD camera. Real-time image videomicroscopy analysis determines the spatial coordinates of the beads with nm accuracy in the x, y and z directions (Pastrana et al., 2016). A step-motor located above the sample moves a pair of magnets allowing the application of stretching forces to the bead-DNA system. Applied forces can be quantified from the Brownian excursions of the bead and the extension of the DNA tether (Strick et al., 1998). Data were acquired at 120 Hz to minimize sampling artefacts in force determination. We used vertically aligned magnets coupled to an iron holder to achieve a force of up to 5 pN.

DNA was diluted and mixed in ParB-Mg^2+^ buffer (50 mM Tris-HCl (pH 7.5), 100 mM NaCl, 0.2 mg/ml BSA, 0.1 % Tween-20, 1 mM DTT, 4 mM MgCl_2_) or ParB-EDTA buffer (by replacing the 4 mM MgCl_2_ with 1 mM EDTA) and then incubated with 1 μm diameter magnetic beads (Dynabeads, Myone streptavidin, Invitrogen) for 10 min. Magnetic beads were previously washed three times and diluted in PBS. DNA:beads ratios were adjusted for each substrate to obtain single tethered beads. Following DNA-beads incubation, then sample was injected in a double parafilm® (Sigma)-layer flow cell. After 10-min adsorption of the beads to the surface we applied a force of 4 pN to remove the non-attached beads and washed with buffer to clean the chamber. Torsionally-constrained molecules and beads with more than a single DNA molecule were identified from its distinct rotation-extension curves (Gutierrez-Escribano et al., 2020) and discarded for further analysis.

Time-course data was obtained by recording the extension of the tether at a low force of 0.33 pN, after a 2-min incubation of DNA tethers and ParB at 4 pN. To obtain force-extension curves, we measured the extension of the tethers at decreasing forces from 5 pN to 0.01 pN for a total measuring time of around 15 min. Force-extension curves were first measured on naked DNA (no ParB data) and always from high to low force. Then, the experiment was repeated on the same molecule but at quoted ParB concentrations. This method allowed us to obtain force-extension curves in the absence and presence of protein for each tethered-DNA molecule. No ParB DNA curves were fitted to the worm-like chain model using Origin Software. Molecules with a discrepancy of contour length of ±15% from the crystallographic length were discarded for the analysis.

### C-trap fluorescence experiments

We used a dual-optical tweezers setup combined with confocal microscopy and microfluidics (C-trap) from Lumicks (Lumicks B.V). Our system has three laser lines (488 nm, 532 nm 535 nm) for confocal microscopy and provides a force resolution of <0.1 pN at 100 Hz, distance resolution of <0.3 nm at 100 Hz, and confocal scanning with < 1 nm spot positioning accuracy (Lumicks). In this work we used a 488 nm laser for illumination and a 500-525 nm filter for its fluorescence. We used a 5 channel microfluidic chamber (**Figure 1 –figure supplement 1A**). Channel 1 contained 4.38 µm SPHERO Streptavidin coated polystyrene beads diluted at 0.005% w/v in fishing buffer (FB, 10 mM Tris-HCl pH 8.0, and 50 mM NaCl). Channel 2 included the 39x *parS* DNA in FB and channel 3 only FB. First, two beads were trapped using the dual optical tweezers in channel 1 and moved to channel 2 to attach DNA molecules to the beads. The capture of DNA was detected by performing a preliminary force-extension curve in channel 2. Then, the bead-DNA system was moved to channel 3, where further force-extension curves were performed to check for single-molecule captures, and to define the zero force point. Finally, a stretching force of 19-23 pN was set, and the bead-DNA system moved to the protein channel 4 that contains ParB_2_^AF488^ at quoted concentrations in ParB-Mg^2+^ buffer supplemented with an oxygen scavenger system (1 mM Trolox, 20 mM glucose, 8 µg/ml glucose oxidase and 20 µg/ml catalase). Confocal images (scans) and kymographs were performed in the protein channel 4 (**Figure 1 –figure supplement 1A**), which was previously passivated with BSA (0.1% w/v in PBS) and Pluronic F128 (0.5% w/v in PBS).

Spreading experiments include a 2 min incubation time in channel 4 before turning on the confocal laser. Spreading-blocking experiments also used four channels but in this case we inject 100 nM EcoRI^E111G^ in ParB buffer in channel 3. Following a 2 min incubation in channel 3 to allow binding of the blocking protein, the bead-DNA system was moved to channel 4, which in this case contained 100 nM EcoRI^E111G^ and 20 nM ParB_2_^AF488^ in ParB buffer supplemented with the oxygen scavenger (see above). An additional 2-min incubation was performed before confocal imaging.

Confocal images of a defined area were taken using a pixel size of 100 nm and a scan velocity of 1 µm/ms. With these parameters, typical images of experiments using single or tandem 39x *parS* DNA were obtained every 2 s and 2.7 s, respectively. Confocal laser intensity at the sample was 3.4 µW.

Kymographs were obtained by single-line scans between the two beads using a pixel size of 100 nm and a pixel time of 0.1 ms. Typical temporal resolution of kymographs taken on single or tandem 39x *parS* DNA were 25 ms and 32 ms, respectively.

### C-trap data analysis

Data acquisition was carried out using Bluelake, the commercial software included in the Lumicks C-trap. This software provides HDF5 files, which can be processed using Lumicks’ Pylake Python package. We used homemade Python scripts to export the confocal scans or kymographs in ASCII matrix files or in TIFF format. Python scripts can be found at https://github.com/Moreno-HerreroLab/C-TrapDataProcessing. Profiles were obtained from ASCII files using a home-written LabView program. Images of scans or kymographs were produced using the WSxM freeware (Horcas et al., 2007). Animated GIFs were produced using Image J from individual scans saved in TIFF.

## SUPPLEMENTAL INFORMATION

This paper contains supplemental information.

## ACKNOWLEDGEMENTS

We are grateful to Michelle Hawkins (University of York) for supplying the EcoRI E111G variant. Work in the MSD lab was supported by the Wellcome Trust (New Investigator Award 100401/Z/12/Z to MSD) and the BBSRC (South West Biosciences Doctoral Training studentship to GLMF). F.M.-H. acknowledges support from the European Research Council (ERC) under the European Union Horizon 2020 Research and Innovation Program (grant agreement 681299). Work in the Moreno-Herrero laboratory was also supported by Spanish Ministry of Economy and Competitiveness grant BFU2017-83794-P (AEI/FEDER, UE to F.M.-H.) and Comunidad de Madrid grants Tec4-Bio – S2018/NMT-4443 and NanoBioCancer – Y2018/BIO-4747 (to F.M.-H.).

## AUTHOR CONTRIBUTIONS

F. de A. B. carried out all C-trap and MT experiments and performed data analysis. C. A.-R. produced all DNA substrates and carried out NTPase assays. C. A.-R. and G. L. M. F. produced all ParB proteins. S. de B. produced Python scripts for analysis of C-trap data. C. L. P. developed methods for magnetic tweezers measurements. M.S.D. and F.M.-H. conceived the project, wrote, reviewed and edited the manuscript. F. M. –H. wrote the first draft of the manuscript and supervised the project. All authors critically reviewed the manuscript and approved the final version.

## FIGURE CAPTIONS

**Figure 1 – figure supplement 1.**
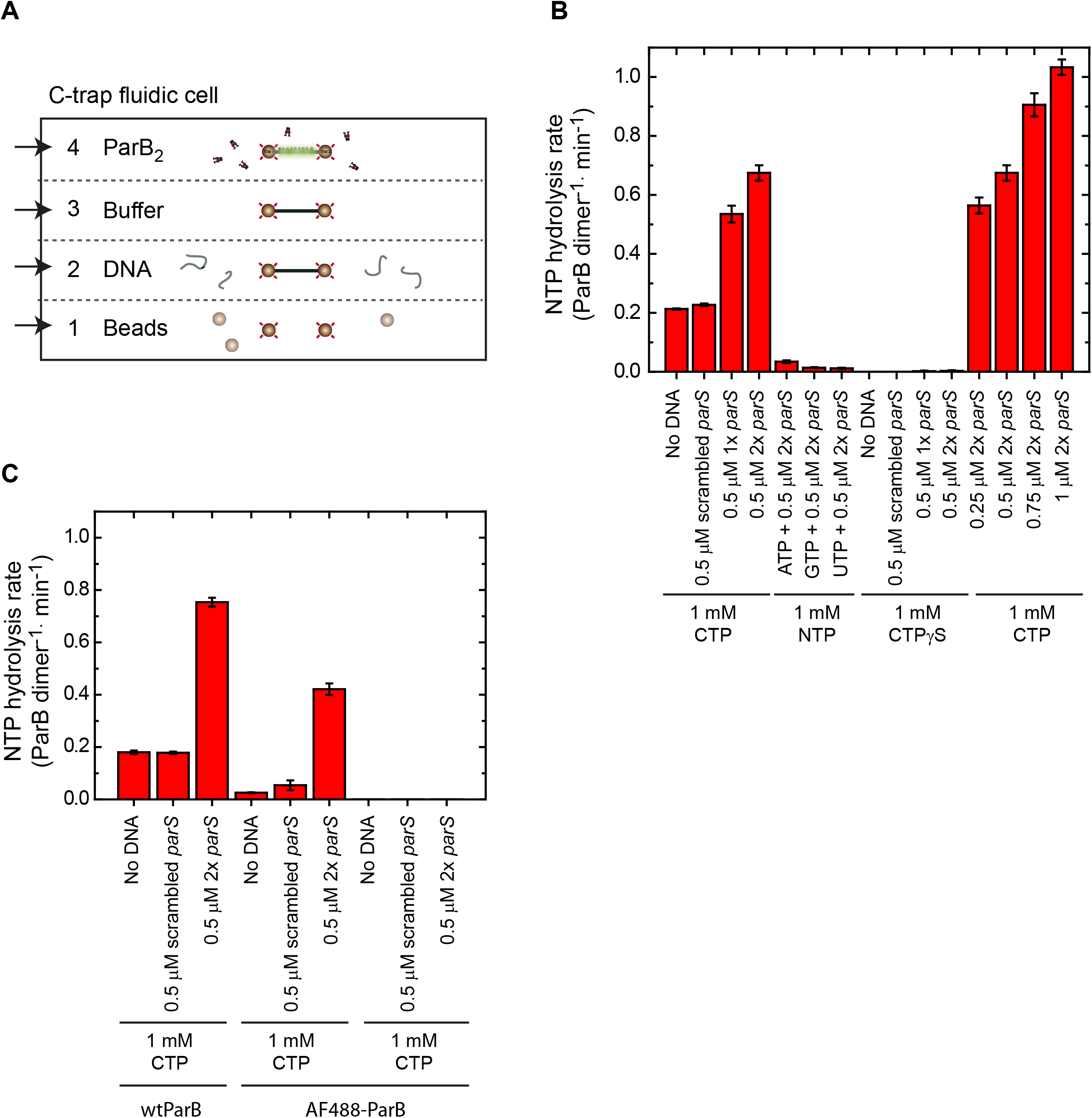
C-trap layout and NTP hydrolysis experiments. (**A**) Cartoon of the fluid chamber employed for the experiments performed in this work. We used four laminar flows, containing streptavidin-coated beads, biotinylated DNA, buffer, or fluorescently labelled proteins. Confocal images and kymographs were obtained in channel 4 after a 2 min protein incubation. (**B**) NTP hydrolysis by wild type ParB was measured using a colorimetric assay. Mean values are shown from four repeat experiments alongside the standard error of the mean. ParB hydrolyses CTP in the absence of DNA at ∼0.2 CTP molecules per dimer, per minute. CTP hydrolysis increases 3-4-fold in the presence of *parS* DNA. Nucleotides ATP, CTPγS, GTP, or UTP cannot be hydrolysed by ParB. The CTP hydrolysis rate increases linearly with the concentration of *parS* sequences. **C**) NTP hydrolysis by wild type ParB or ParB_2_^AF488^ was measured in the absence or presence of DNA. Mean values are shown from two repeat experiments alongside the standard error of the mean. ParB^AF488^ retains *parS*-stimulated CTPase activity within 2-fold levels of wild type protein.

**Figure 1 – figure supplement 2.**
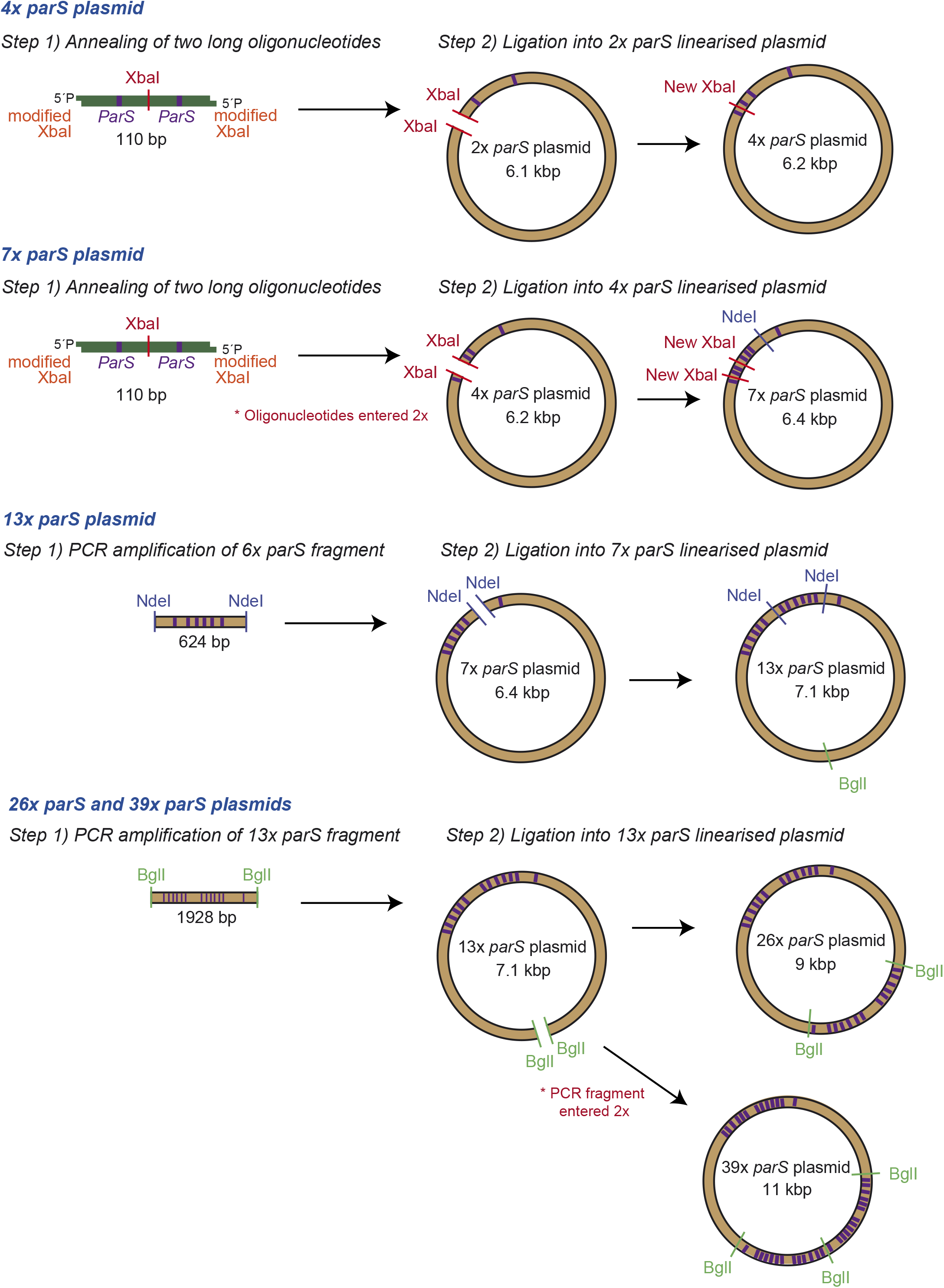
Fabrication of small DNA plasmids. Schematic representation of the steps followed to fabricate DNA plasmids containing multiple copies of *parS* which were used to prepare the magnetic tweezers DNA substrates.

**Figure 1 – figure supplement 3.**
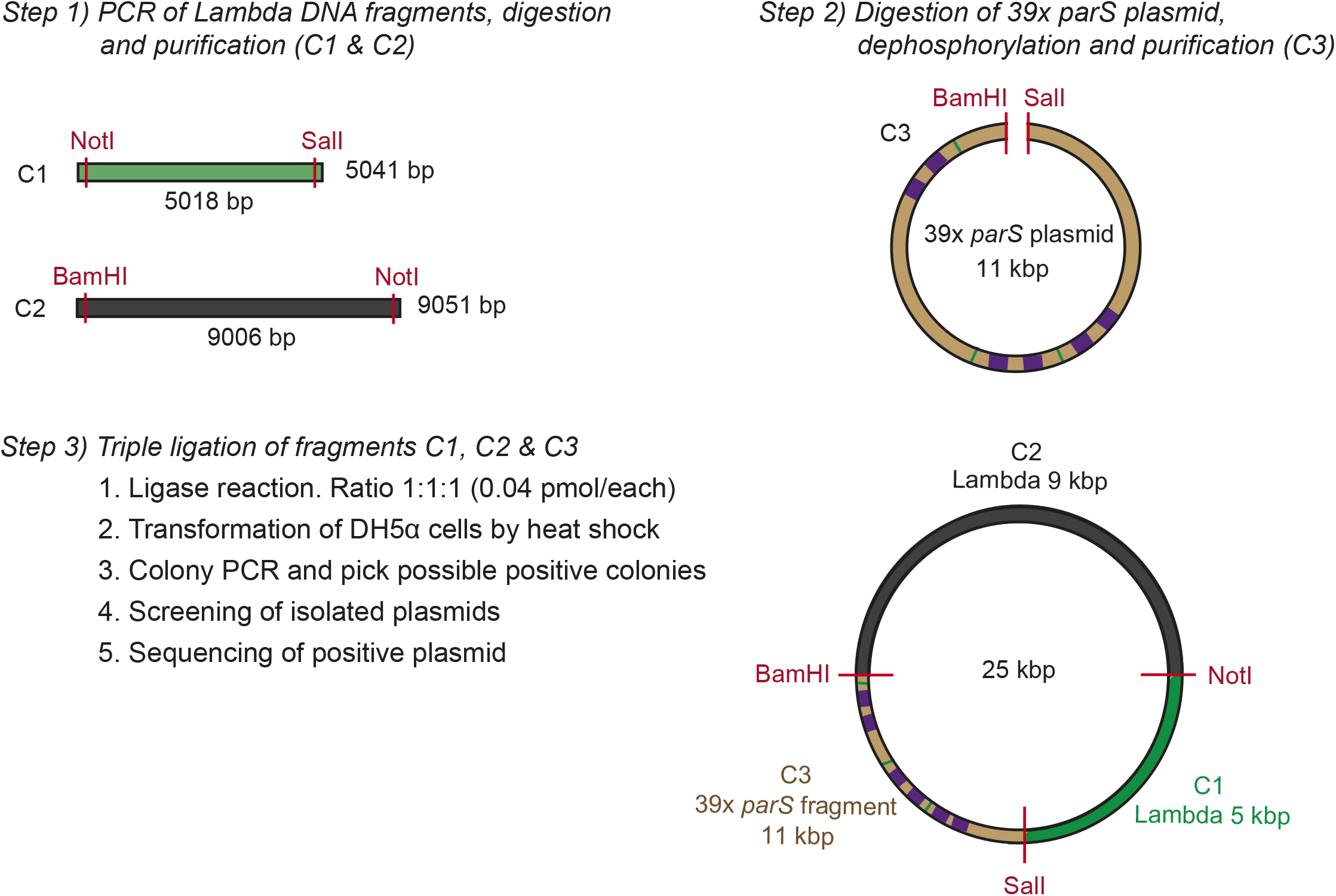
Fabrication of large DNA plasmids. Schematic representation of the steps followed to fabricate the large plasmids used to make the C-trap DNA substrates. This scheme specifically represents the cloning of the large 39x *parS* plasmid. Further details are described in the **Supplementary Methods** section.

**Figure 1 – figure supplement 4.**
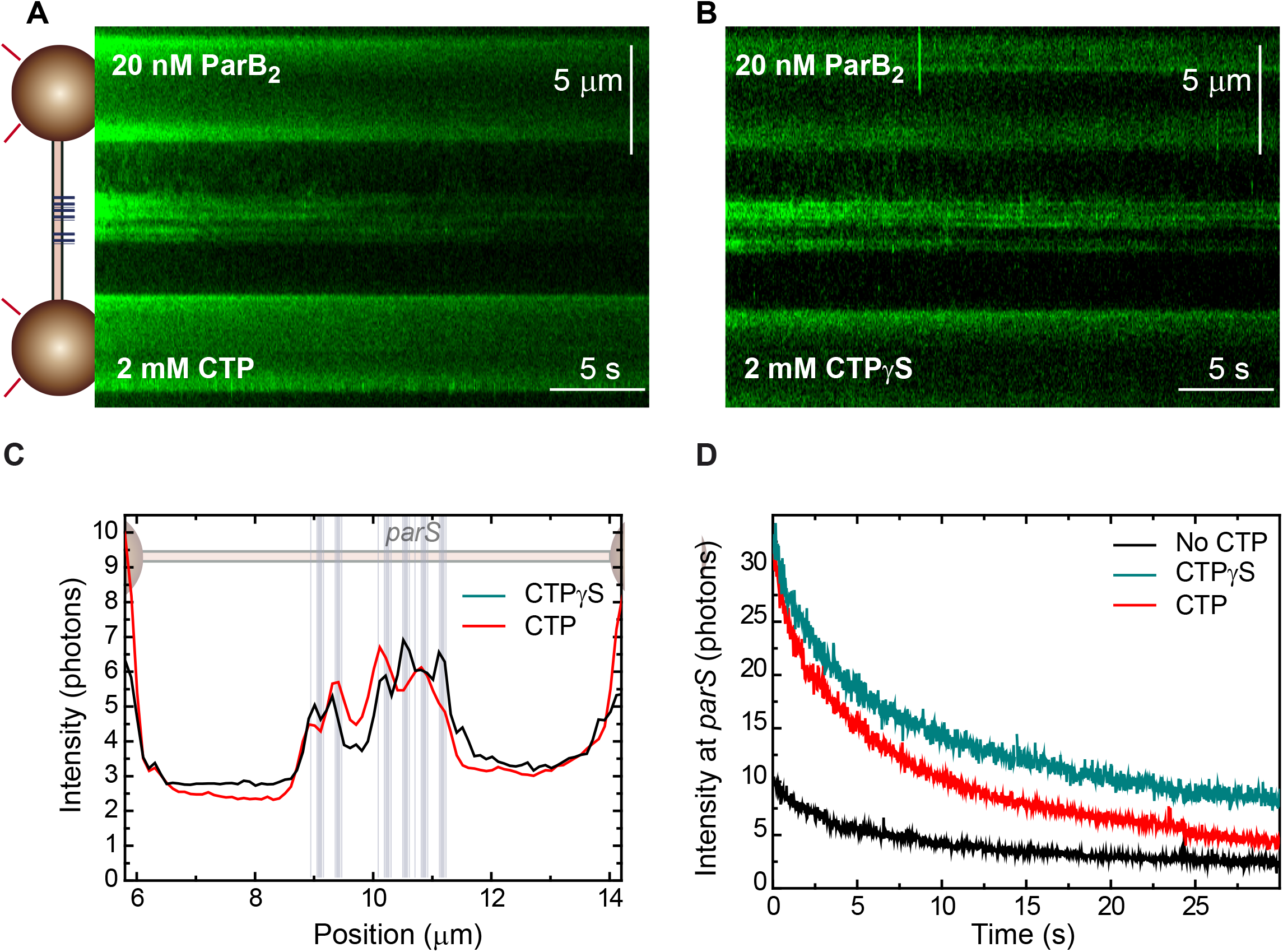
(**A**) Fluorescence kymograph of a single 39x *parS* DNA molecule (**Figure 1C**) obtained with the C-trap including 20 nM ParB2 and 2 mM CTP. A cartoon of the experiment with a schematic of the DNA showing the positions of the *parS* sequences plotted to scale is included on the left. (**B**) Same experiment as in **A** but using 2 mM CTPγS. Regions corresponding to the position of the *parS* groups could be clearly identified. (**C**) Average intensity profile (30 s) obtained along the DNA molecule. Positions of the *parS* sequences are included to scale in the background. The fluorescence intensity peaks correlate very well with the position of the 6 groups of *parS* sites. (**D**) Time-evolution of the fluorescence intensity at the *parS* region obtained from kymographs under no CTP, CTP, and CTPγS conditions.

**Figure 3 – figure supplement 1.**
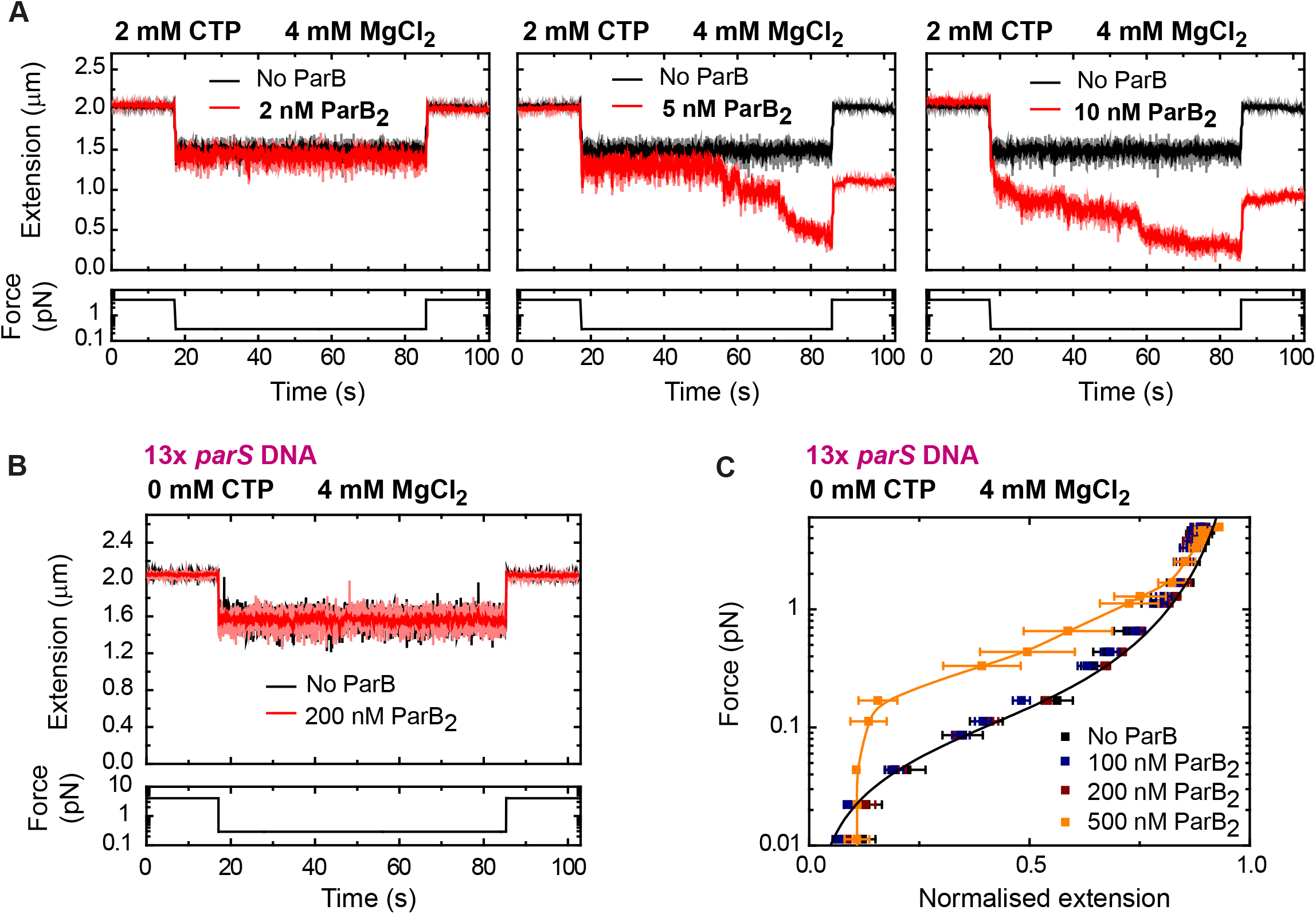
Low nanomolar concentrations of ParB condense parS DNA in the presence of CTP-Mg^2+^. (**A**) Condensation time-courses showing that a minimum of 5 nM ParB2 condenses 13x *parS* DNA within a minute under 2 mM CTP, 4 mM MgCl_2_ conditions. (**B**) Control experiment showing no condensation in the absence of CTP. (**C**) A protein concentration above 500 nM condenses DNA via non-specific interactions, as previously reported (Taylor et al., 2015). No ParB data represent force-extension curves of DNA taken in the absence of protein and are fitted to the worm-like chain model. Errors are standard error of the mean for measurements taken on different molecules (N ≥ 7).

## SUPPLEMENTARY METHODS

### Fabrication of DNA plasmids with multiple copies of *parS*

All constructs were cloned into the vector pET28a(+) (Novagen). Plasmids were cloned in DH5α Competent cells (ThermoFisher Scientific), and after selection of possible positive colonies by colony PCR, plasmids were purified from cultures using QIAprep Spin Miniprep Kit (QIAGEN), analyzed by restriction digestion and finally checked by DNA sequence analysis. These plasmids were employed to prepare different magnetic tweezers substrates which sequences are included below.

#### ‘Scrambled’ parS DNA

The cloning of a plasmid containing a ‘scrambled’ *parS* site (scrambled *parS*: 5’-**C**GT**G**CC**CA**G**G**GA**G**ACA) was described in (Taylor et al., 2015).

#### 1x parS DNA

The cloning of a plasmid containing a 1x *parS* sequence (optimal sequence of *B. subtilis parS* = 5’-TGTTCCACGTGAAACA) was also described in (Taylor et al., 2015).

#### 2x parS DNA

The plasmid containing 2x *parS* sequences was derived from the 1 x *parS* plasmid by ligation of annealed synthetic oligonucleotides containing the *parS* sequence with appropriate overhangs into the cut NcoI site.

#### 4x parS DNA

Two long 5’-phosphorylated oligonucleotides (**Table S1**) containing 2x *parS* sites separated by an XbaI restriction site and ending in a modified XbaI restriction site, were hybridised as described in **Materials and Methods** section. The plasmid containing 2x *parS* sequences was digested with XbaI, dephosphorylated with rSAP (NEB) and ligated with this pair of hybridized oligonucleotides rendering a plasmid with 4x *parS* sites.

The oligonucleotides were designed to create an incomplete XbaI site at both ends after ligation in such a way that once ligated, XbaI cannot cut again in those positions. In addition, in the middle of the annealed oligonucleotides a bona-fide XbaI site was included, allowing the repetition of the ligation process to insert the annealed oligonucleotides as many times as desired to increase the number of copies of the *parS* site (**Figure 1-figure supplement 2)**.

#### 7x parS DNA

The plasmid containing 4x *parS* sequences was digested with XbaI, dephosphorylated and ligated again with the pair of hybridized oligonucleotides rendering a plasmid with 6x *parS* sites (not employed in this paper). However, in this step of cloning and by chance, the pair of hybridized oligonucleotides entered two times during the ligation process. Although that ligation should render a plasmid with 8x *parS* sites, notice that one of the *parS* sites was incomplete (TTTCACGTGGAACA) probably because an error during the synthesis of one of the oligonucleotides, and therefore that incomplete sequence was not considered as a *parS* site and the final construct is said to contain 7x *parS* sequences.

#### 13x parS DNA

The plasmid containing 7x *parS* sequences was used as a template to amplify by PCR (Phusion Polymerase, NEB) the 6 *parS* fragment by using primers 32.F pET28 PCR NdeI and 33.R pET28 PCR NdeI (**Table S1**). The PCR fragment was digested with NdeI, and ligated into the plasmid with 7x *parS* sequences previously digested with NdeI and dephosphorylated, rendering a plasmid with 13x *parS* sites

#### 26x parS DNA and 39x parS DNA

The plasmid containing 13x *parS* sequences was used as a template to amplify by PCR the 13 *parS* fragment by using 50.F pET28 37 SphI-BglI and 51.R pET28 BglI-SphI (**Table S1**). The PCR fragment was digested with BglI, and ligated into the plasmid with 13x *parS* sequences previously digested with BglI and dephosphorylated, rendering a plasmid with 26x *parS* sites. In addition, in this step of cloning and by chance, in a different plasmid the PCR fragment entered two times during the ligation process, and therefore we also obtained a plasmid with 39x *parS* sites. This last one was employed to prepare large plasmids described below.

### Fabrication of large plasmids (>17 kbp) for C-trap experiments

#### 39x parS DNA

The large plasmid containing 39x *parS* sites was obtained by ligation of three fragments of DNA (**Figure 1 – figure supplement 3**). Two of them correspond to two PCR-fabricated DNAs (C1 and C2) derived from Lambda DNA (NEB) as template and including suitable restriction sites in the designed primers (**Table S1**). Fragment C1 (green), that corresponds to positions 33464-38474 bp of Lambda DNA, was amplified with oligos 20Lambda_F_NotI and 21Lambda_R_SalI, purified and digested with NotI and SalI. Fragment C2 (black), that corresponds to positions 5475-14509 bp of Lambda DNA, was amplified with oligos 22Lambda_F_BamHI and 23Lambda_R_NotI, purified and digested with NotI and BamHI. The fragment containing the 39x *parS* sites (C3, brown in the scheme) was obtained by digestion of the plasmid containing 39x *parS* sites described above with BamHI and SalI, dephosphorylation and purification by gel extraction with Gel extraction Kit (QIAGEN). The three fragments were then ligated at a ratio 1:1:1 and DH5α Competent cells were transformed by regular heat shock procedure. After selection of possible good colonies, large plasmids were purified from cultures using QIAprep Spin Miniprep Kit and checked by restriction digestion followed by DNA sequence analysis.

#### EcoRI 39x parS DNA

The large plasmid containing the 39x *parS* region flanked by two regions of 5xEcoRI restriction sites (named as EcoRI 39x *parS*) was produced following the same procedure but replacing parts C1 and C2 with fragments C1-EcoRI and C2-EcoRI, each one including 5 EcoRI restriction sites at the desired position. However, during the PCR amplification of Fragment C2-EcoRI with oligos 177.Lambda_F_5Eco BamHI and 23Lambda_R_NotI, the forward oligo was annealed in a different position of the Lambda DNA due to a similar sequence, and therefore the amplified fragment was shorter and corresponds to positions 13254-14509 bp of lambda DNA.

**Supplementary Table S1.**
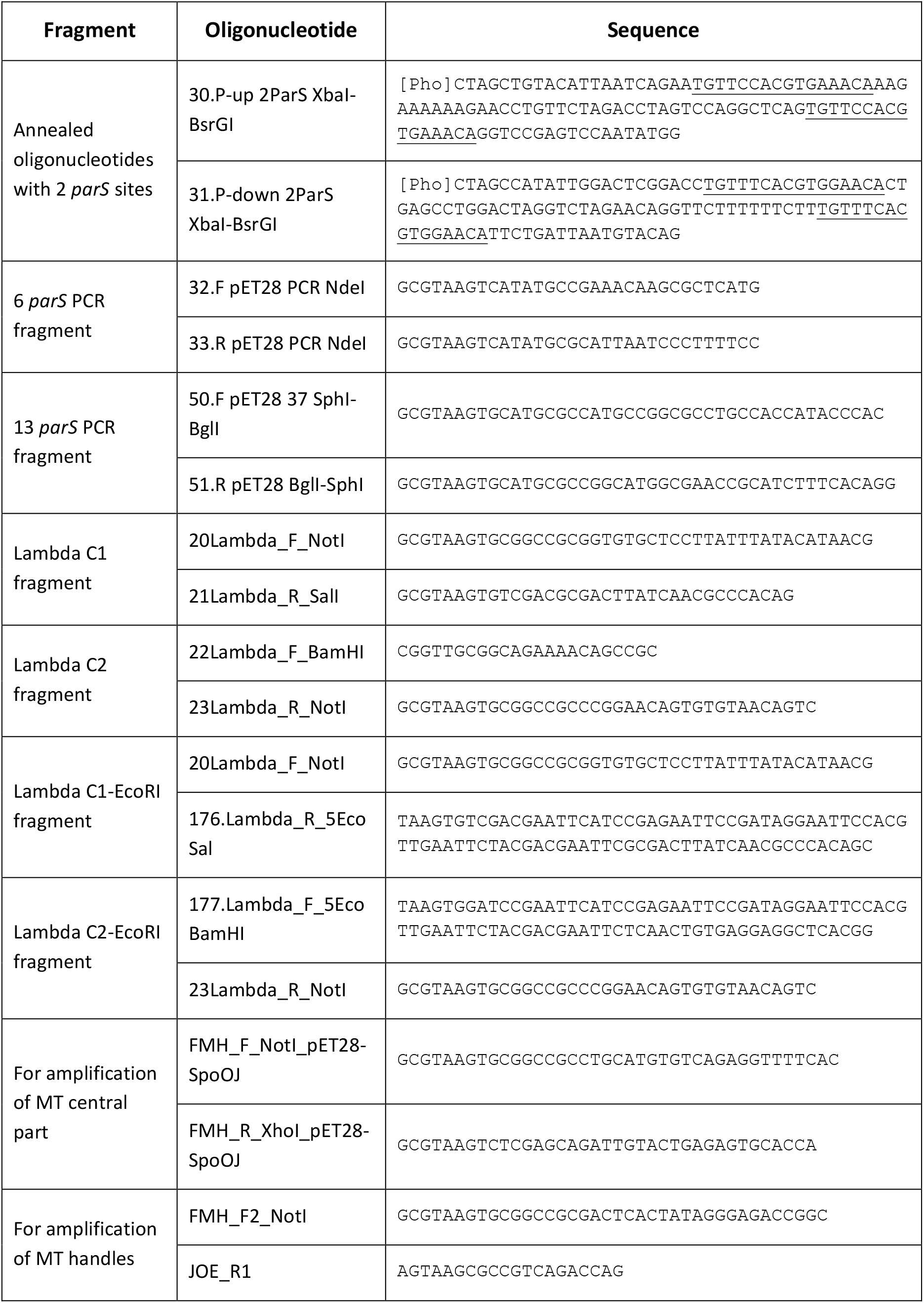
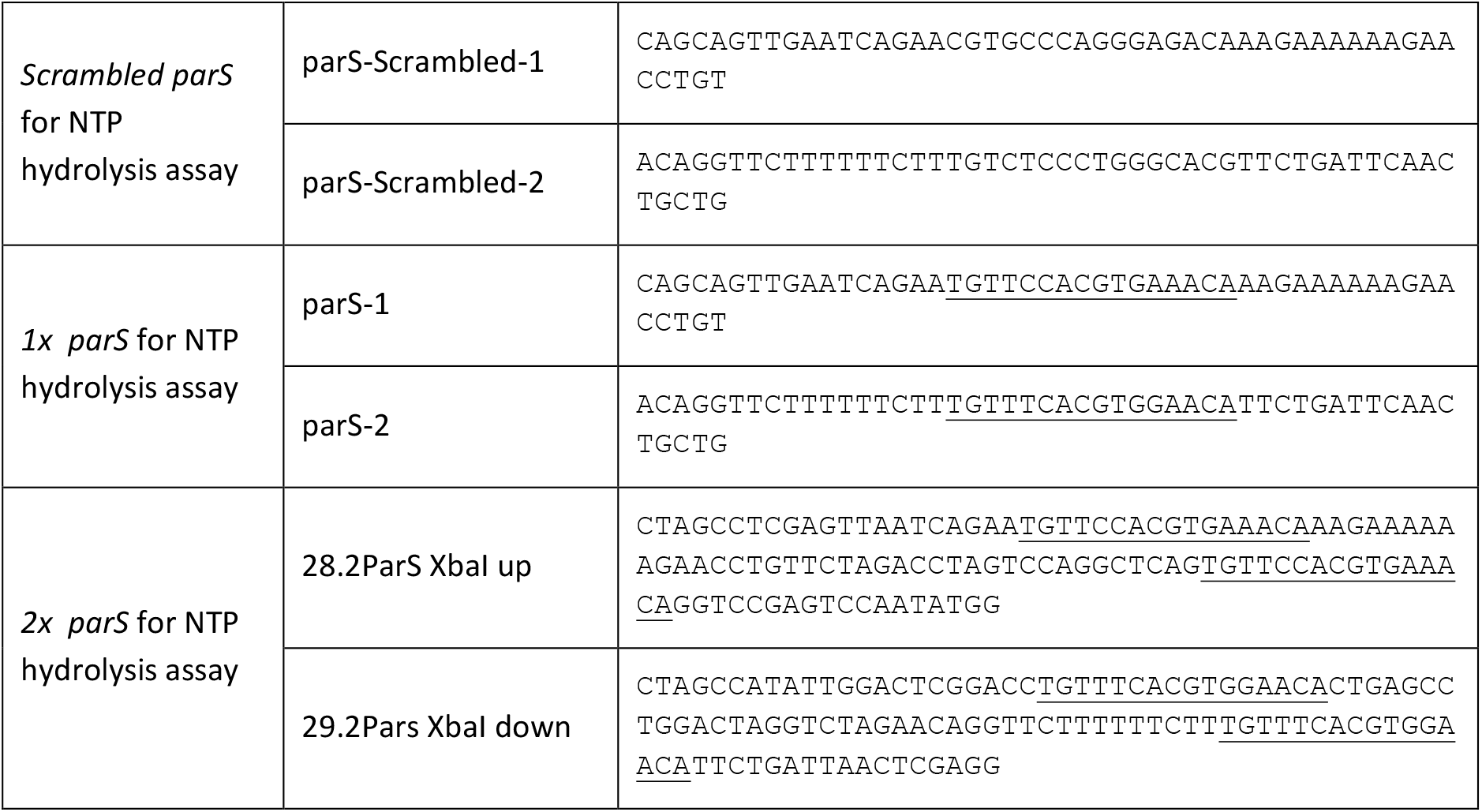
DNA oligonucleotides used in this work.

### Sequences of DNA fragments used in this work

DNA sequences corresponding to the central part (not including the handles) of Magnetic Tweezers and C-trap DNA substrates employed in this work. Scrambled *parS* sequence is highlighted in pink. *ParS* sequences in inverted (TGTTTCACGTGGAACA) or direct (TGTTCCACGTGAAACA) orientation are highlighted in yellow. EcoRI restriction sites present in the central part of the C-trap substrate EcoRI 39x *parS* DNA are highlighted in red.

#### MT ‘Scrambled’ parS DNA (5971 bp)

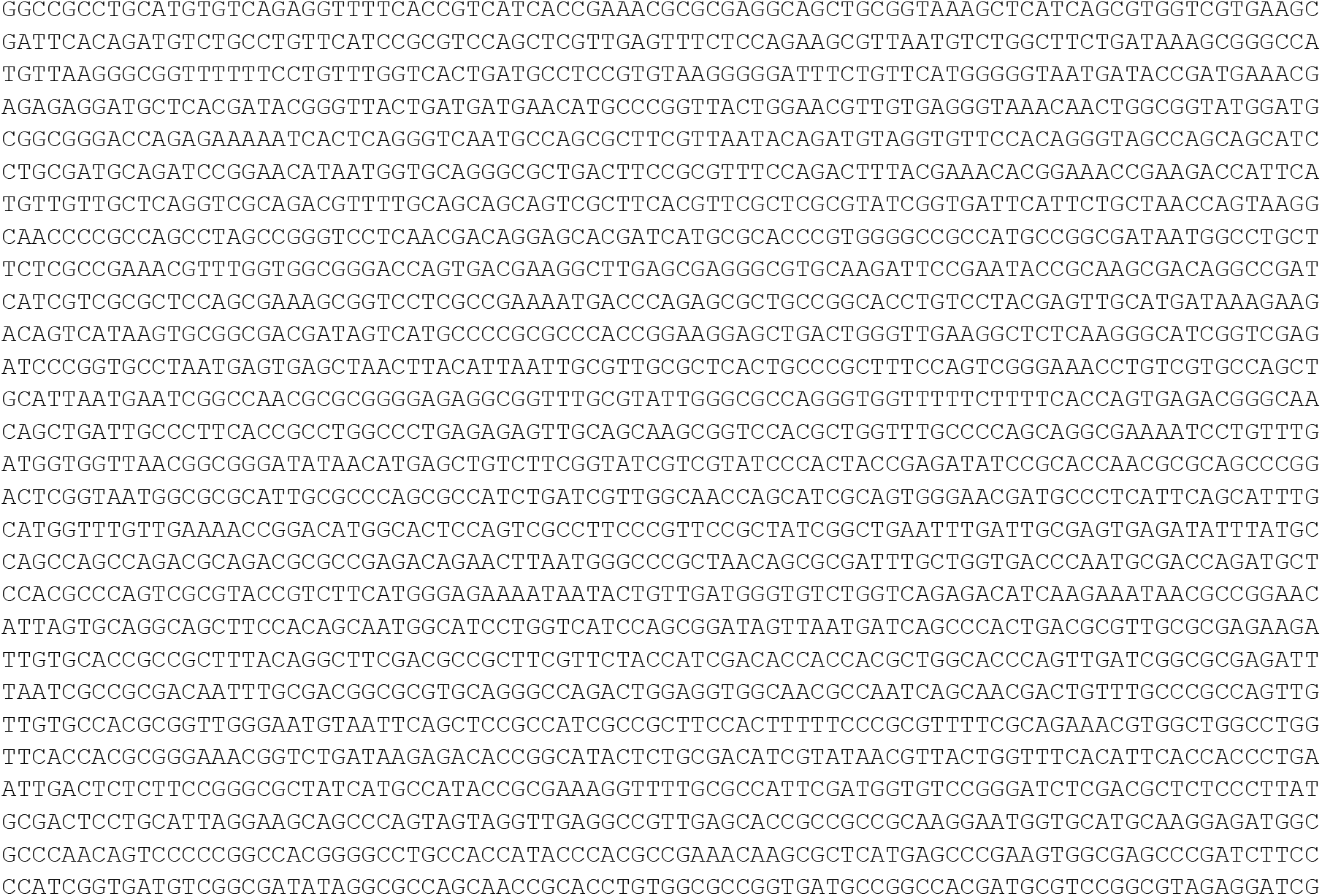
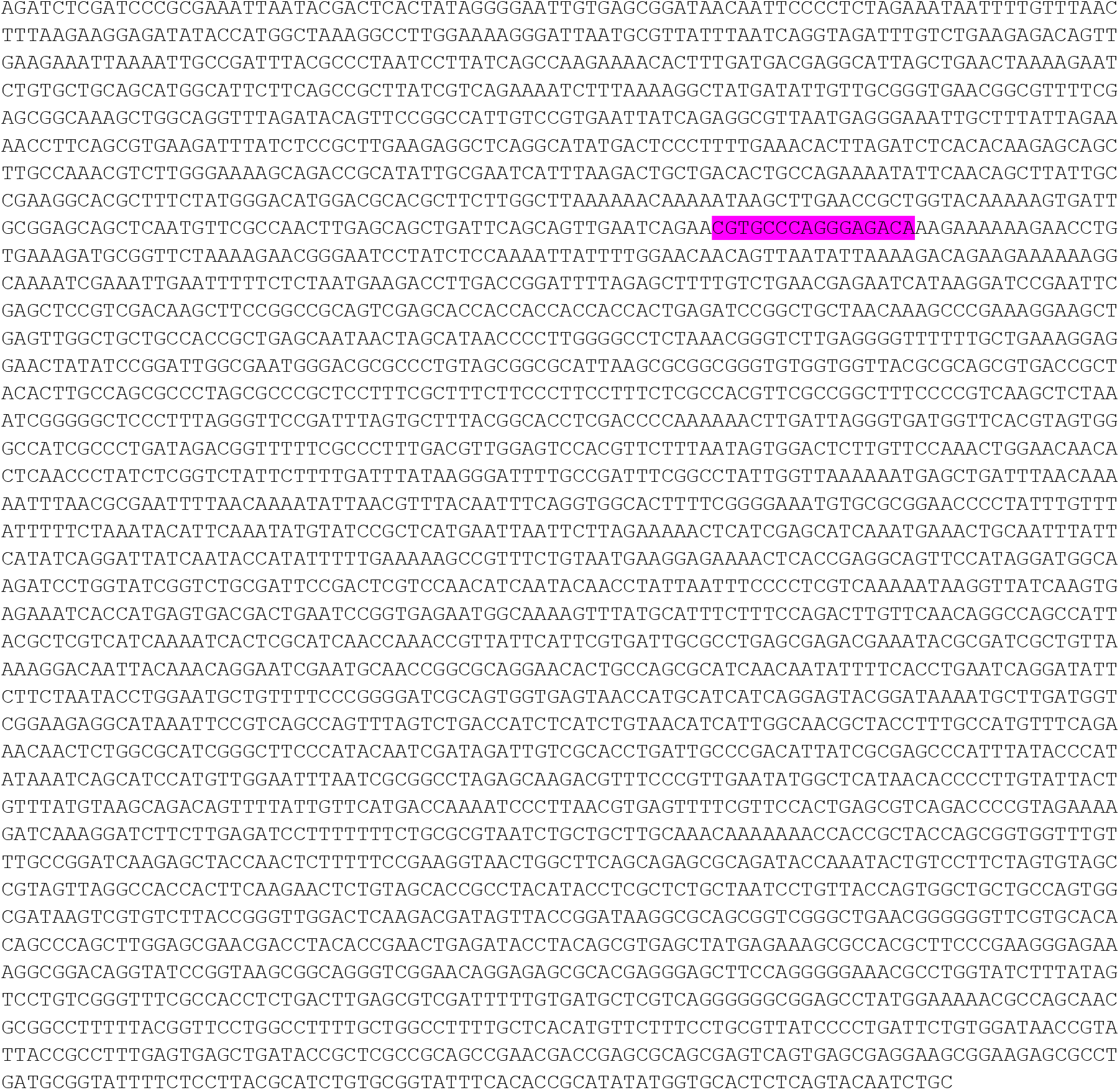

#### MT 1x parS DNA (5971 bp)

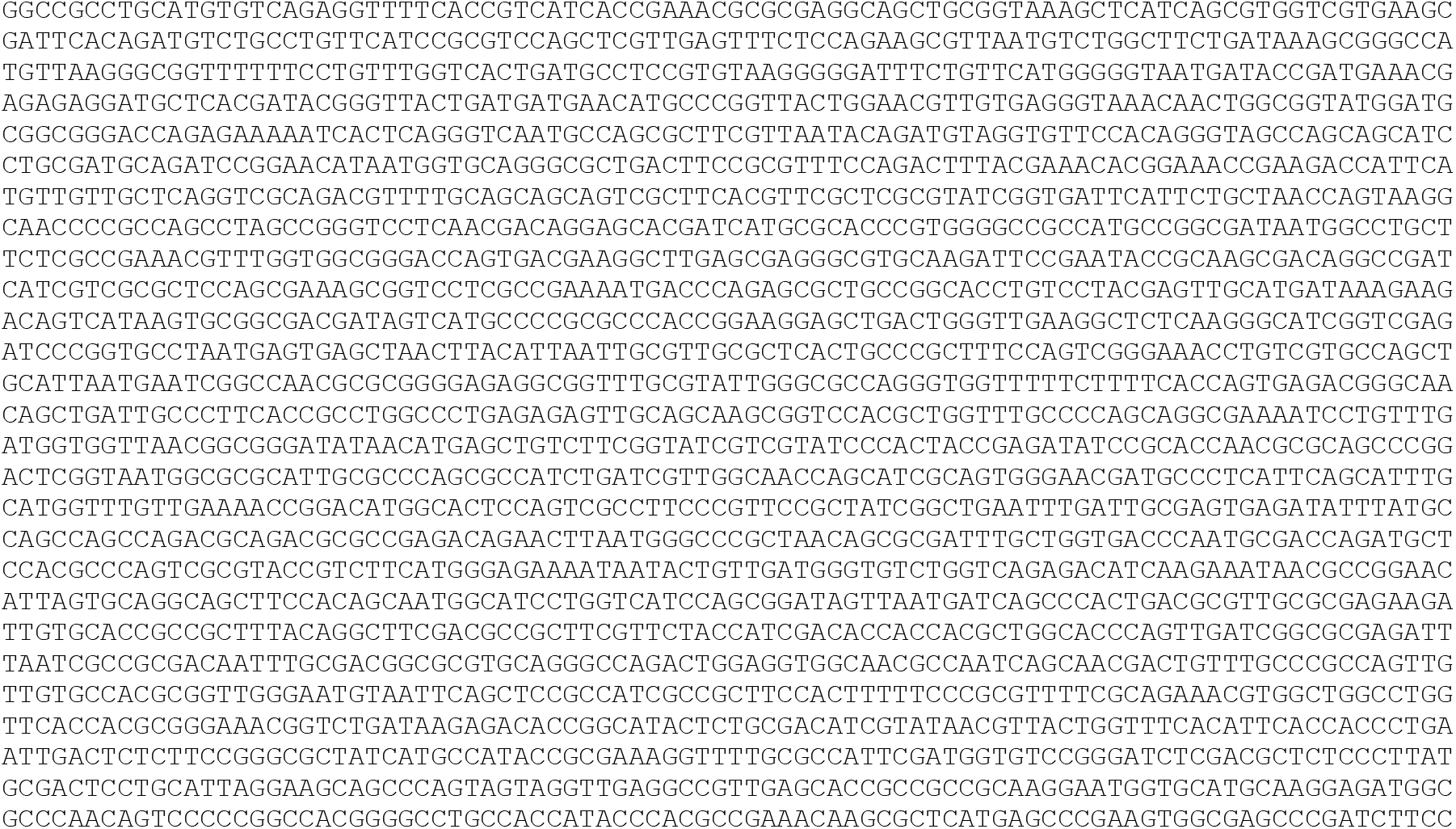
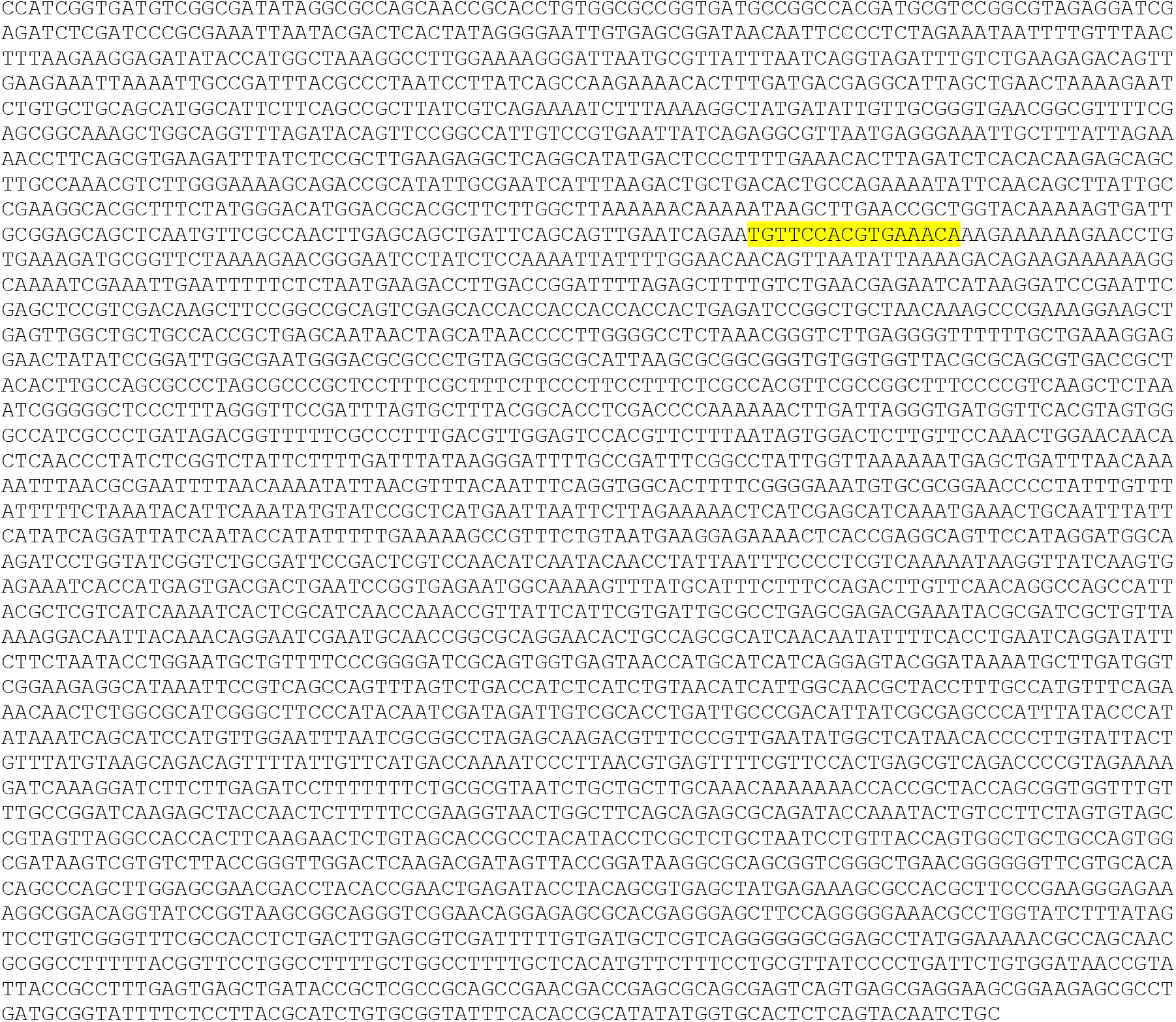

#### MT 2x parS DNA (5991 bp)

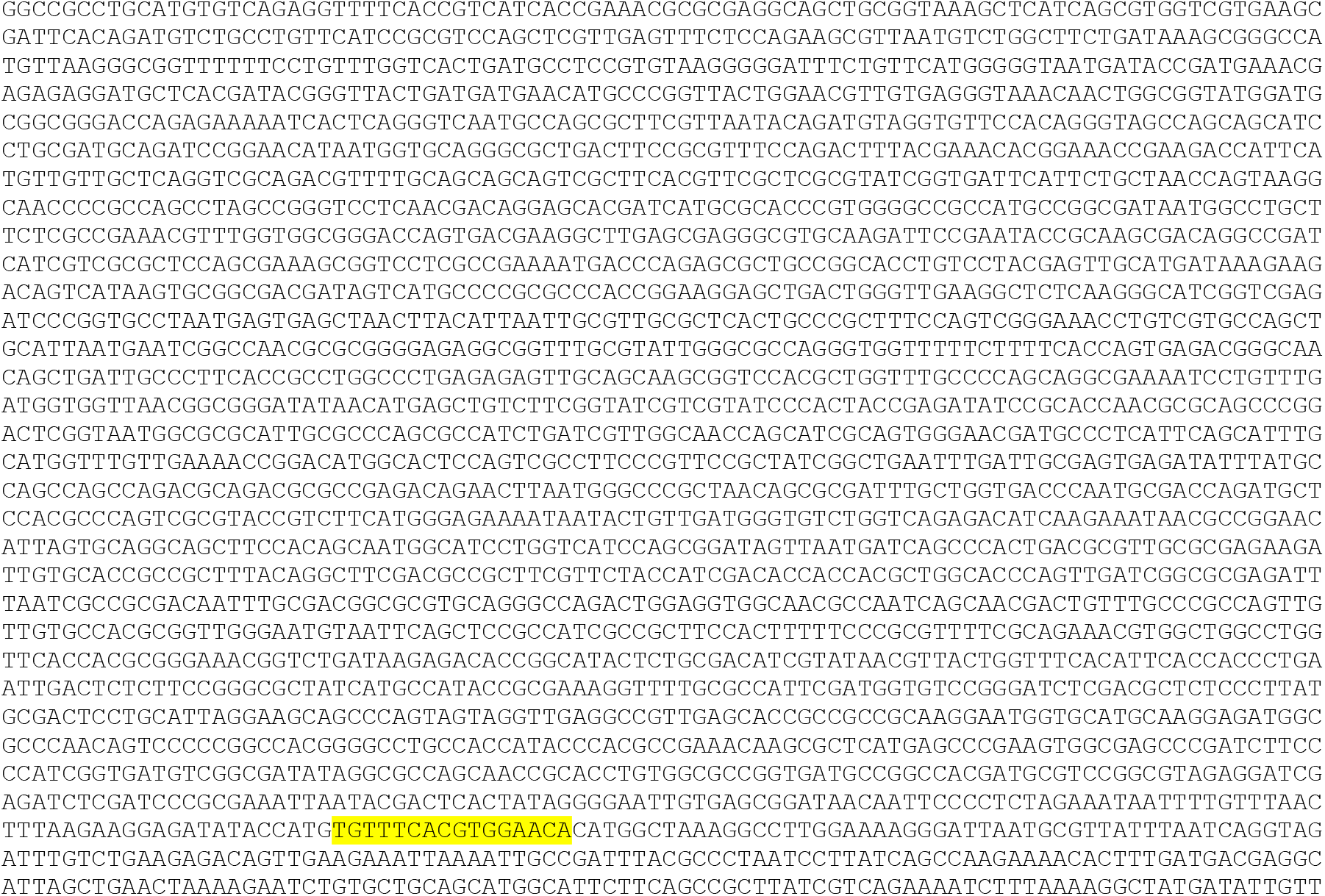
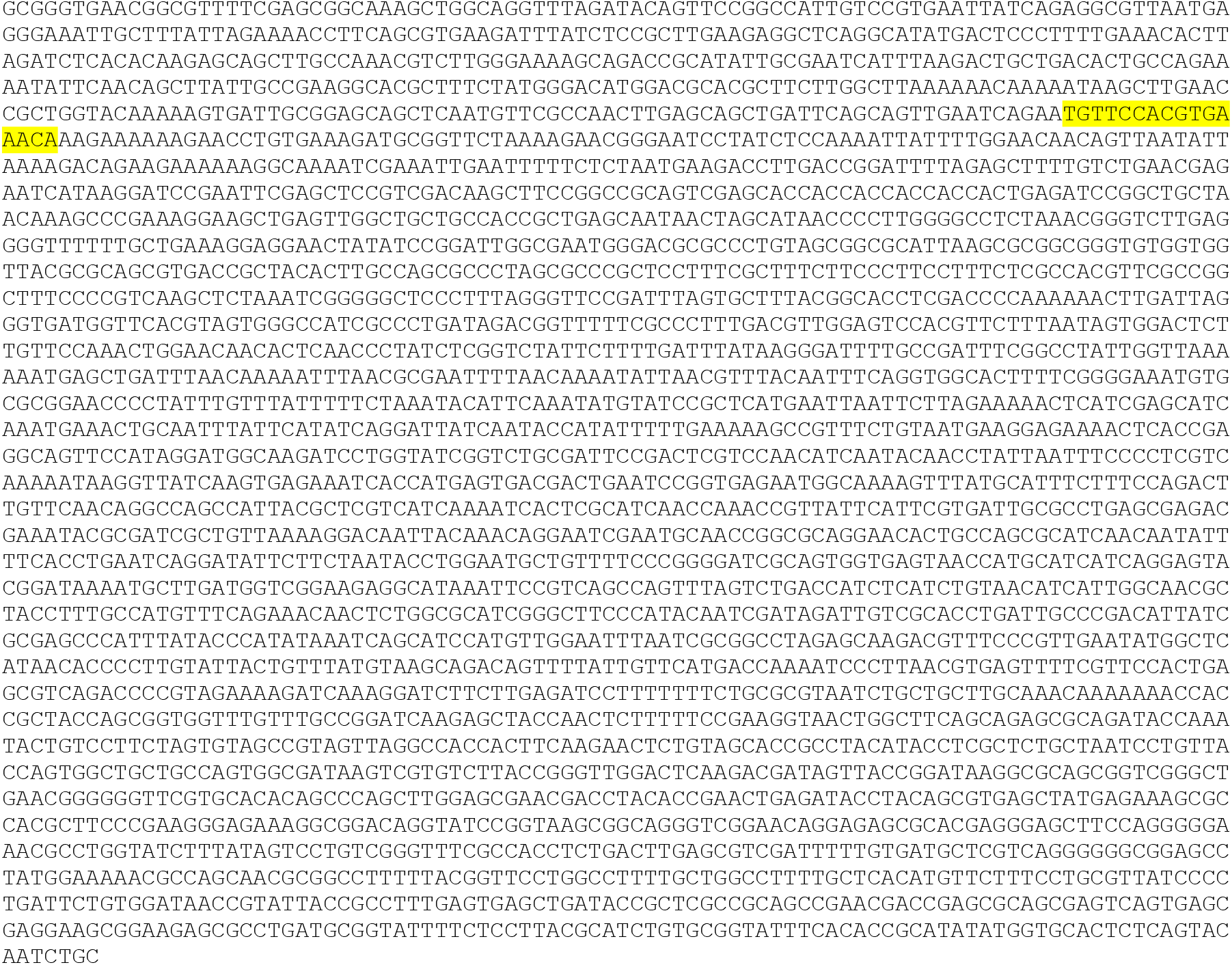

#### MT 4x parS DNA (6101 bp)

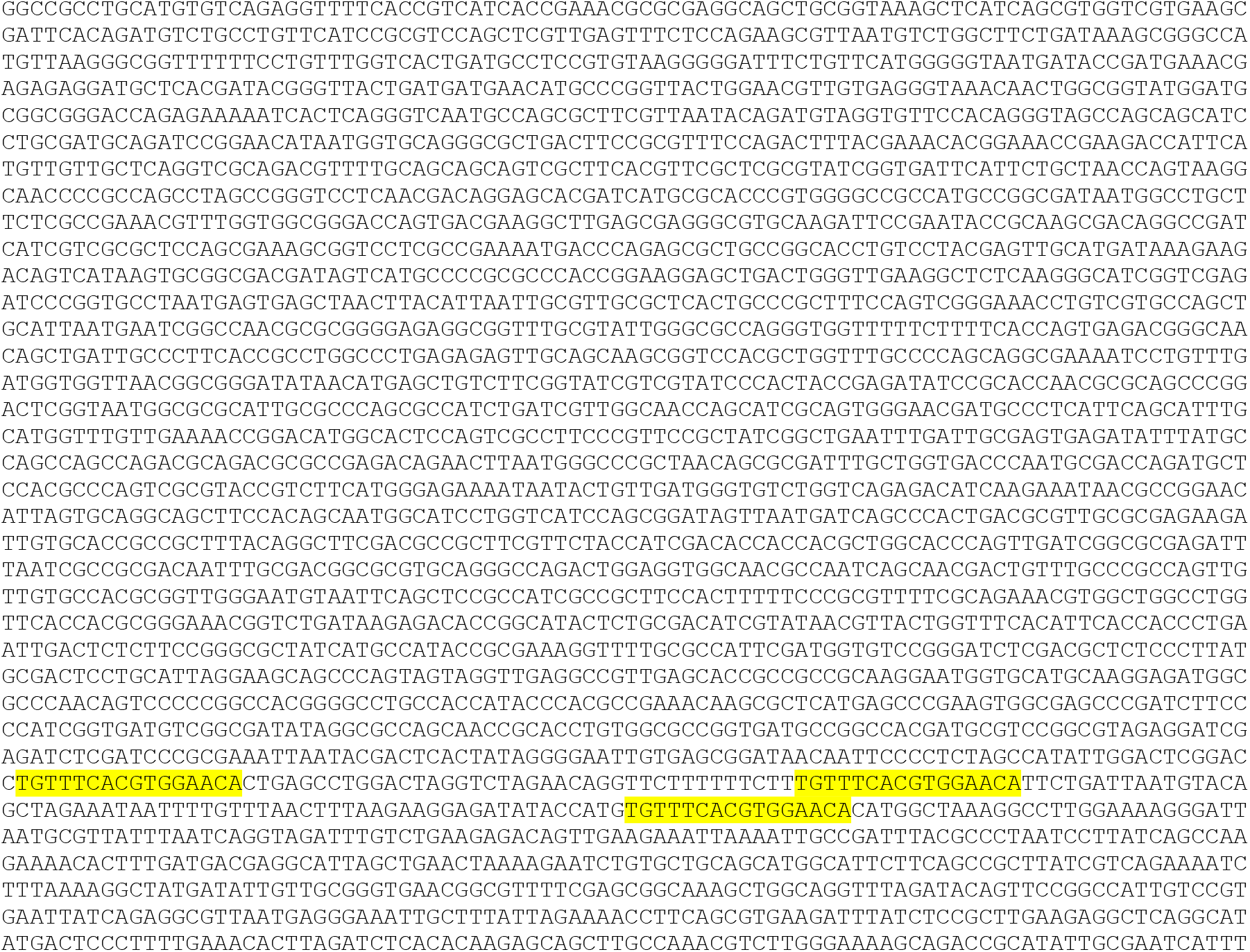
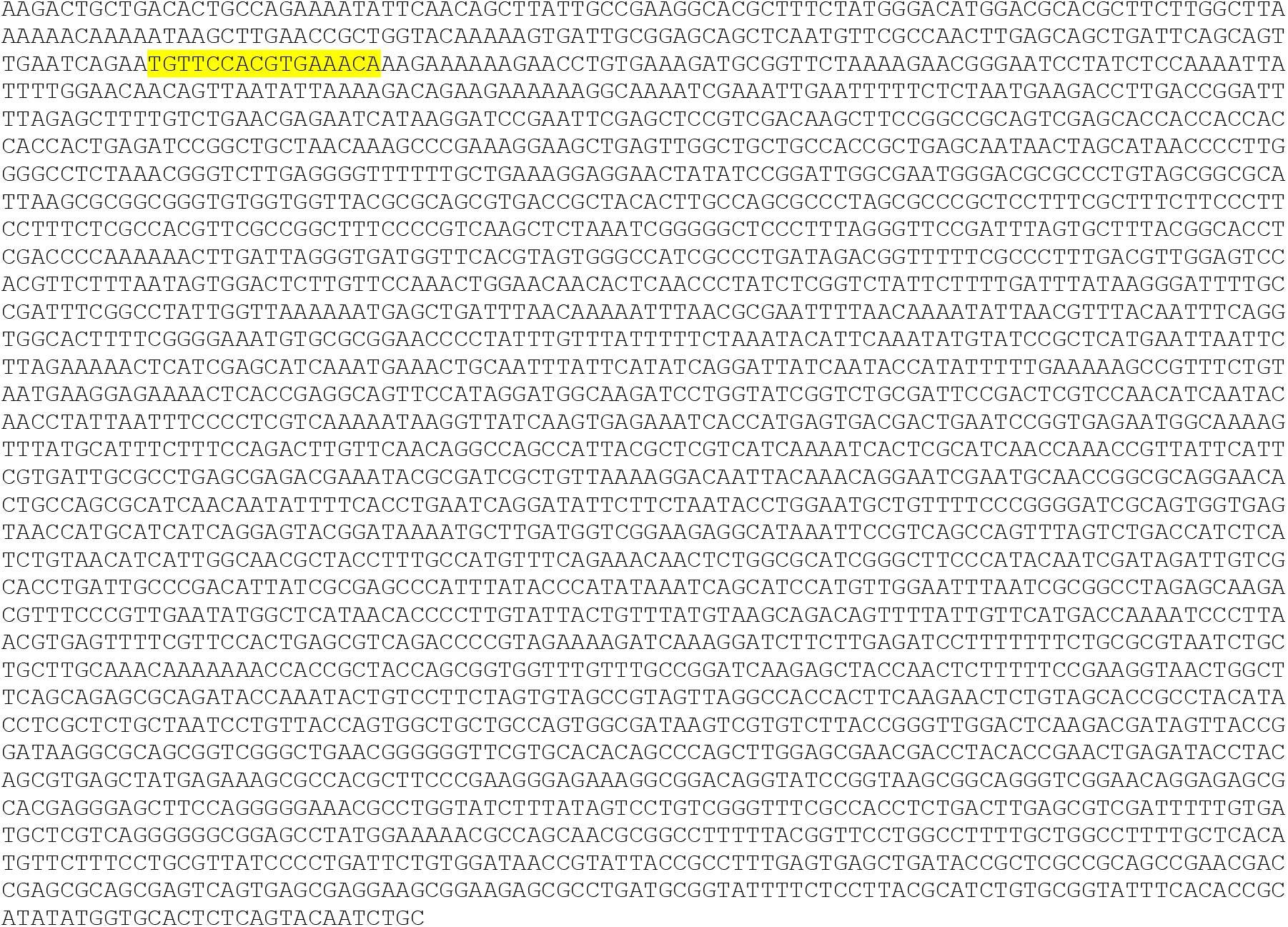

#### MT 7x parS DNA (6316 bp)

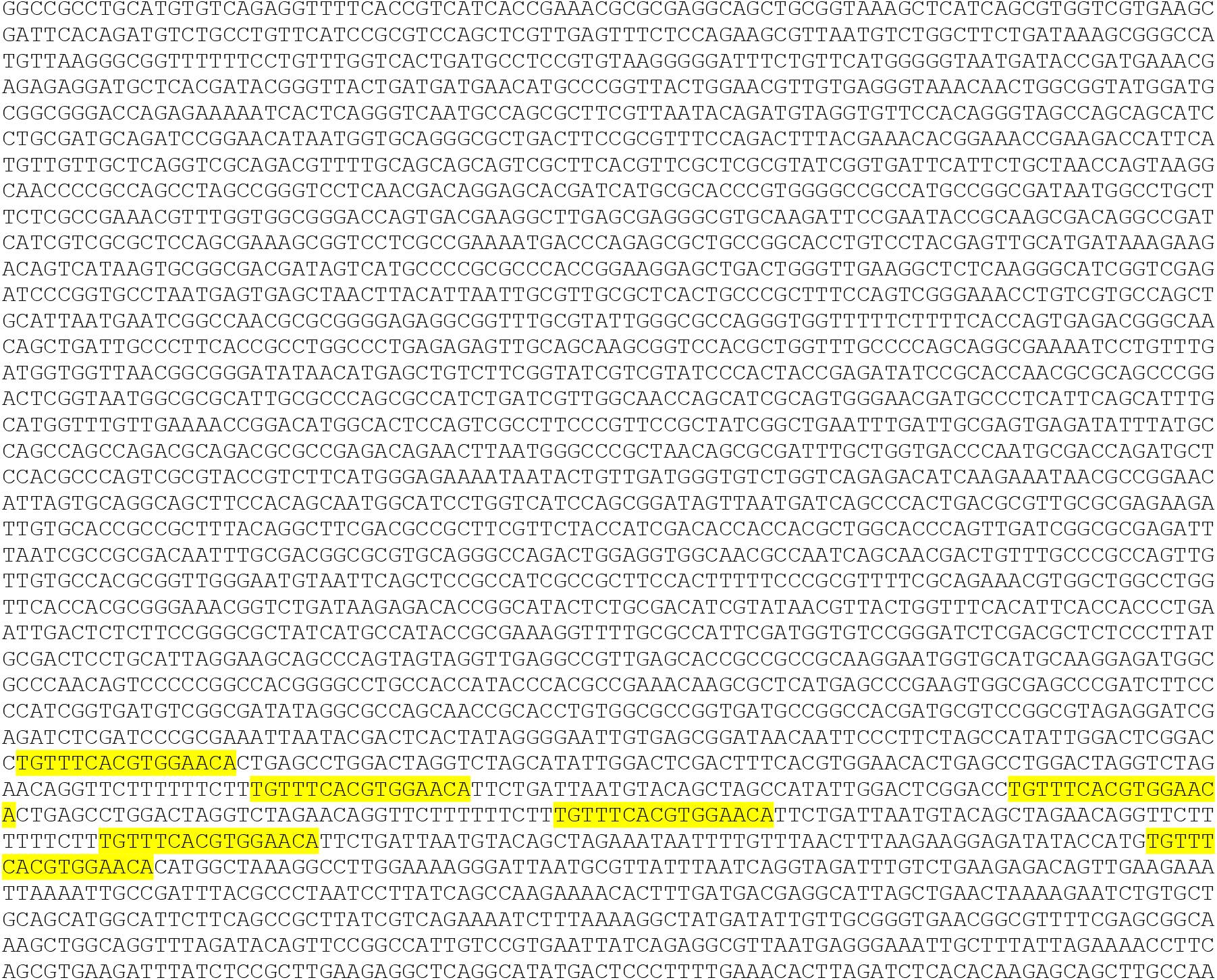
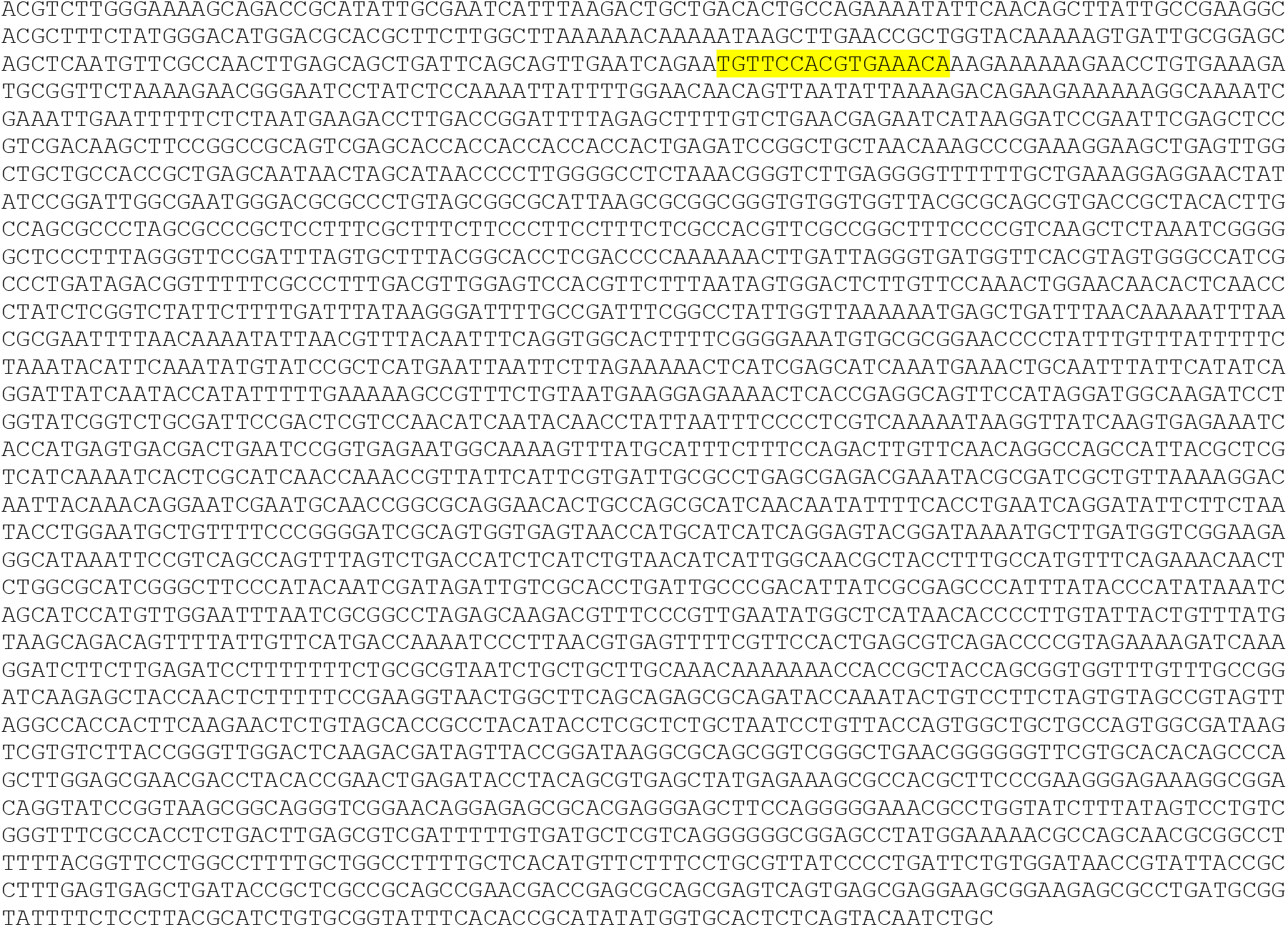

#### MT 13x parS DNA (6940 bp)

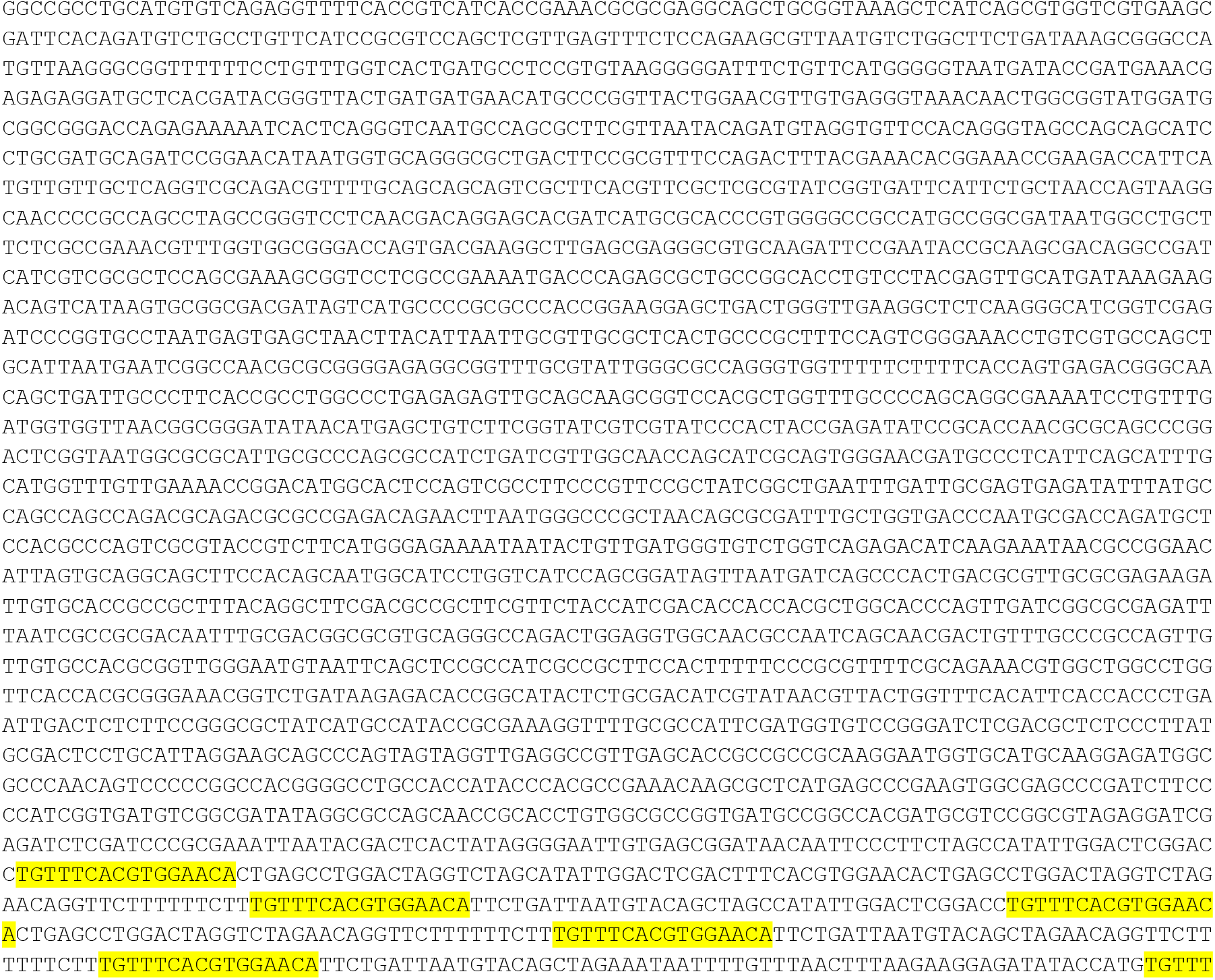
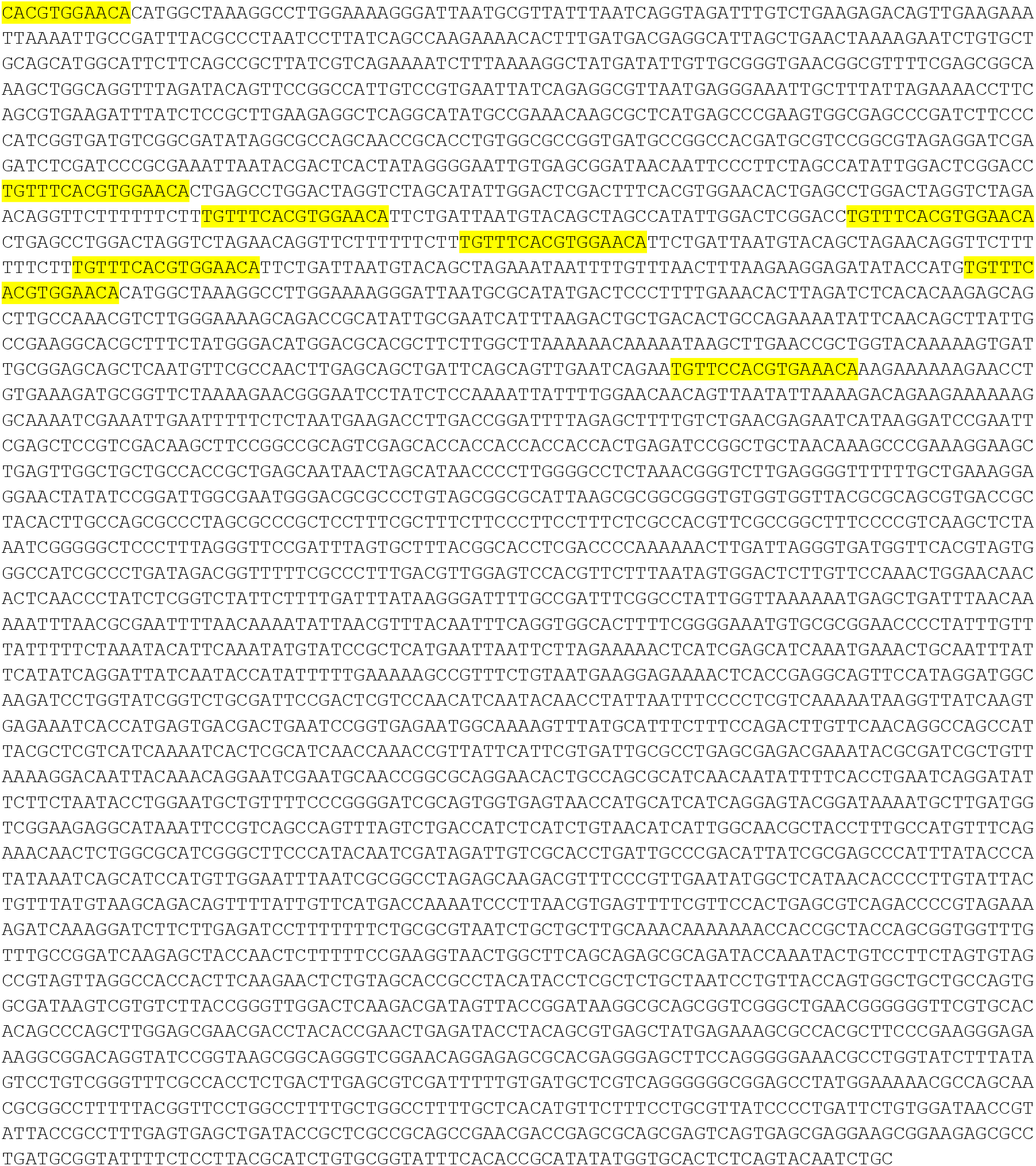

#### MT 26x parS DNA (6348 bp)

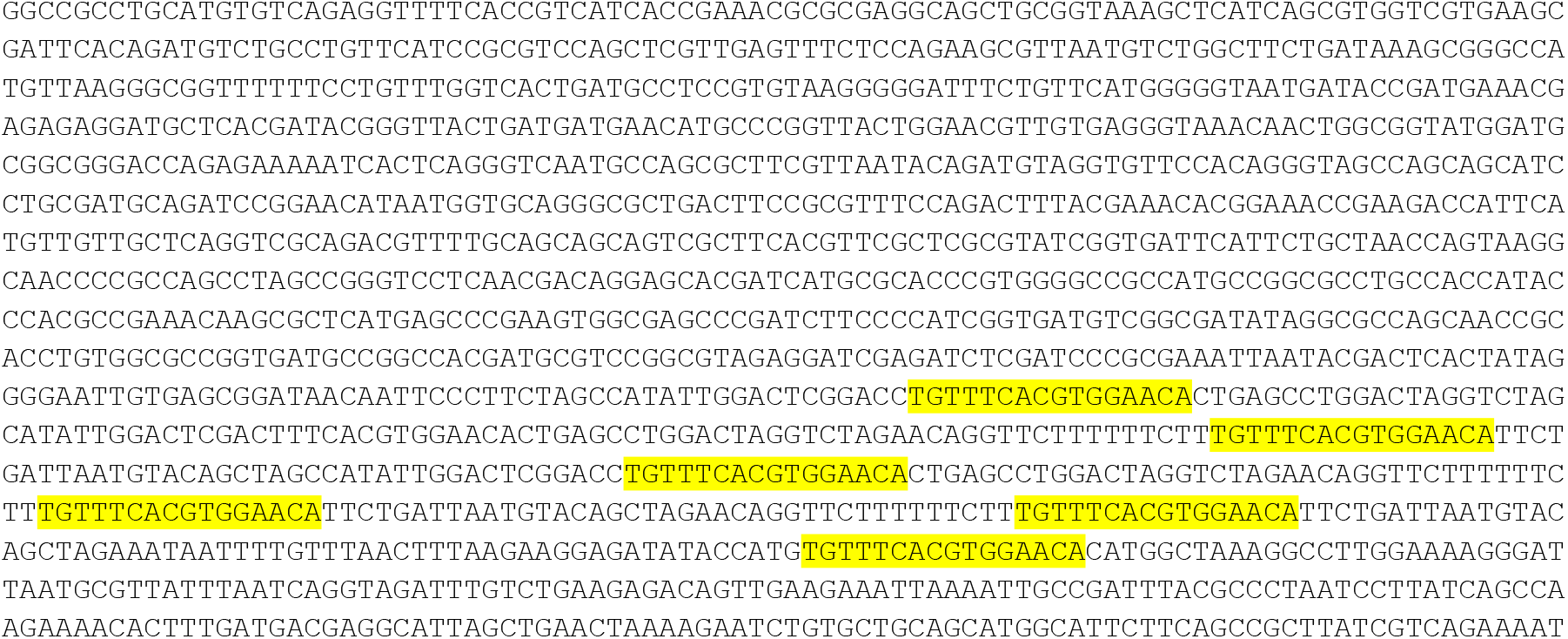
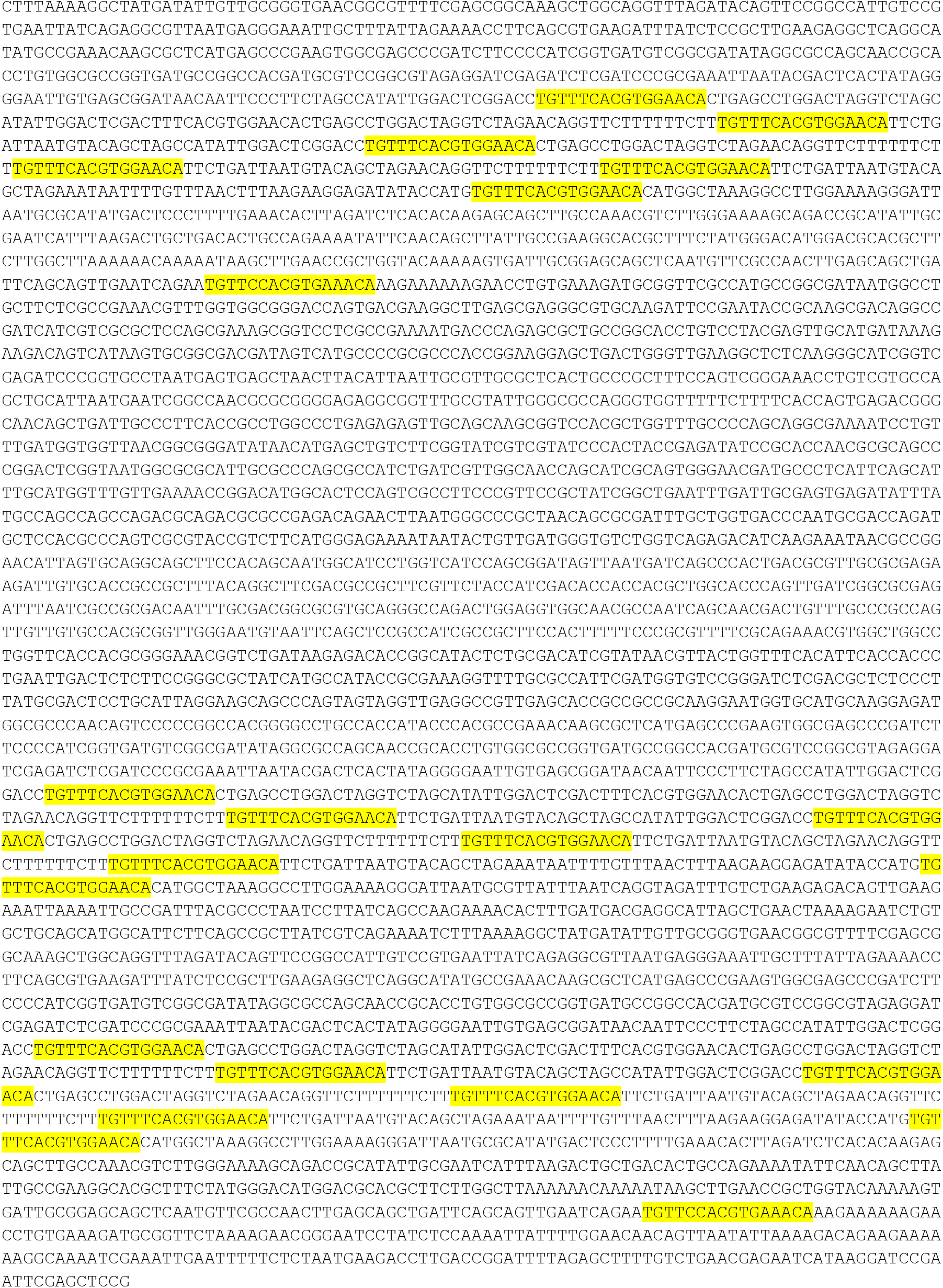

#### C-trap 39x parS DNA (24974 bp)

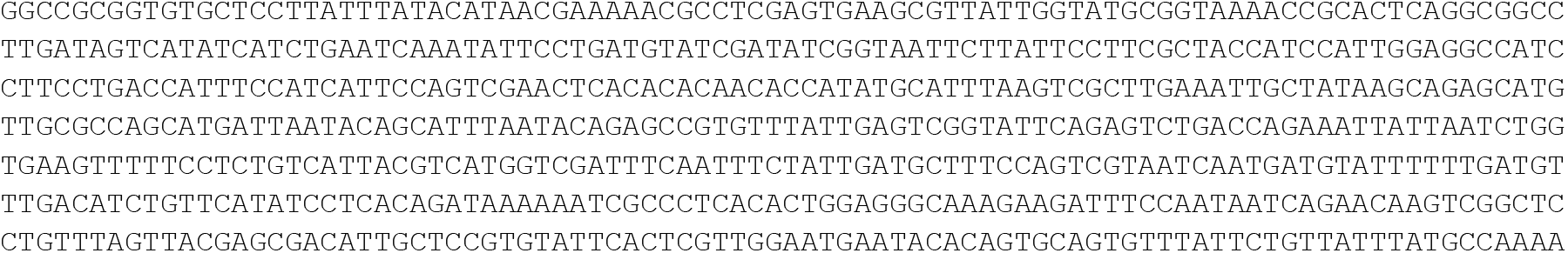
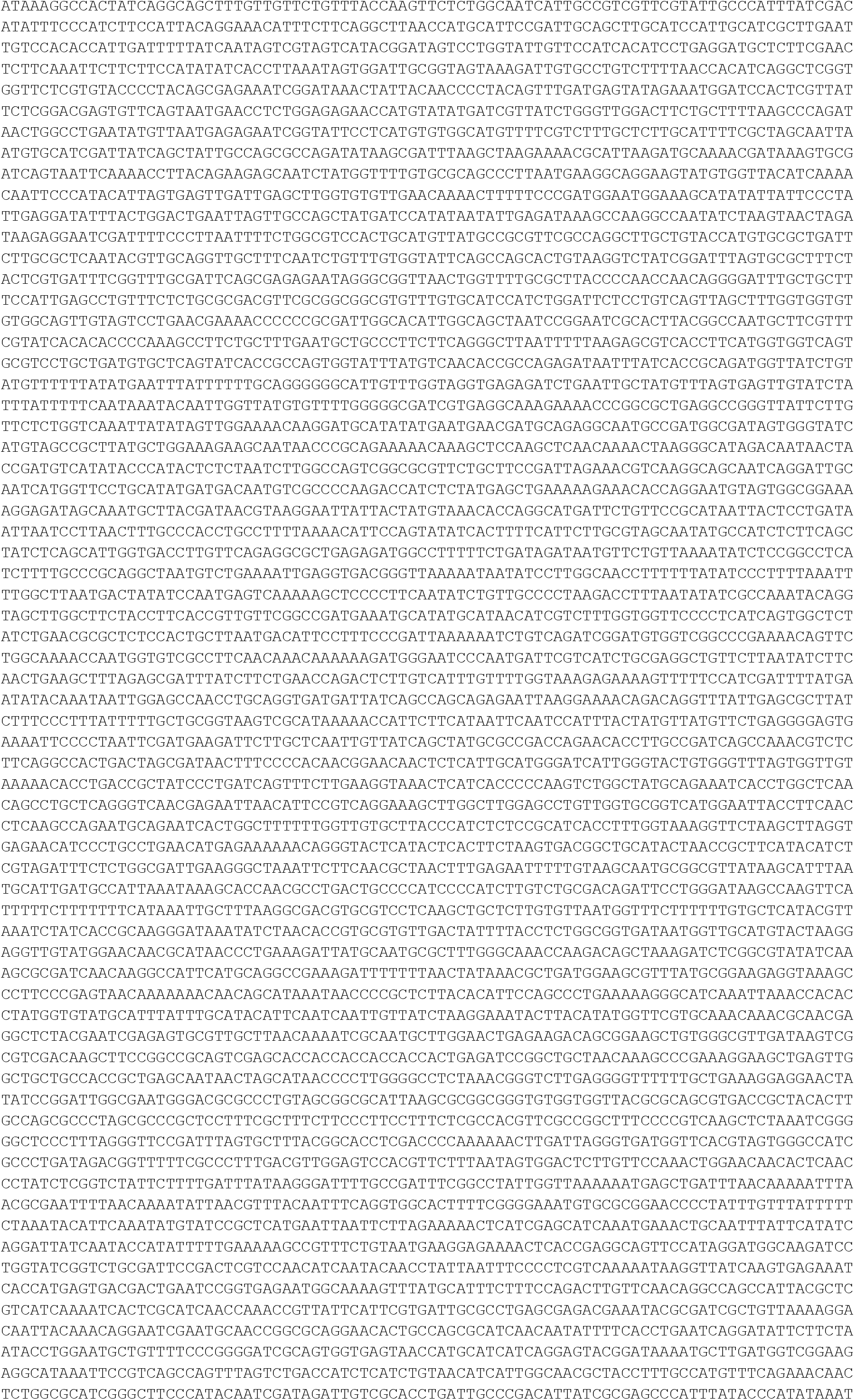
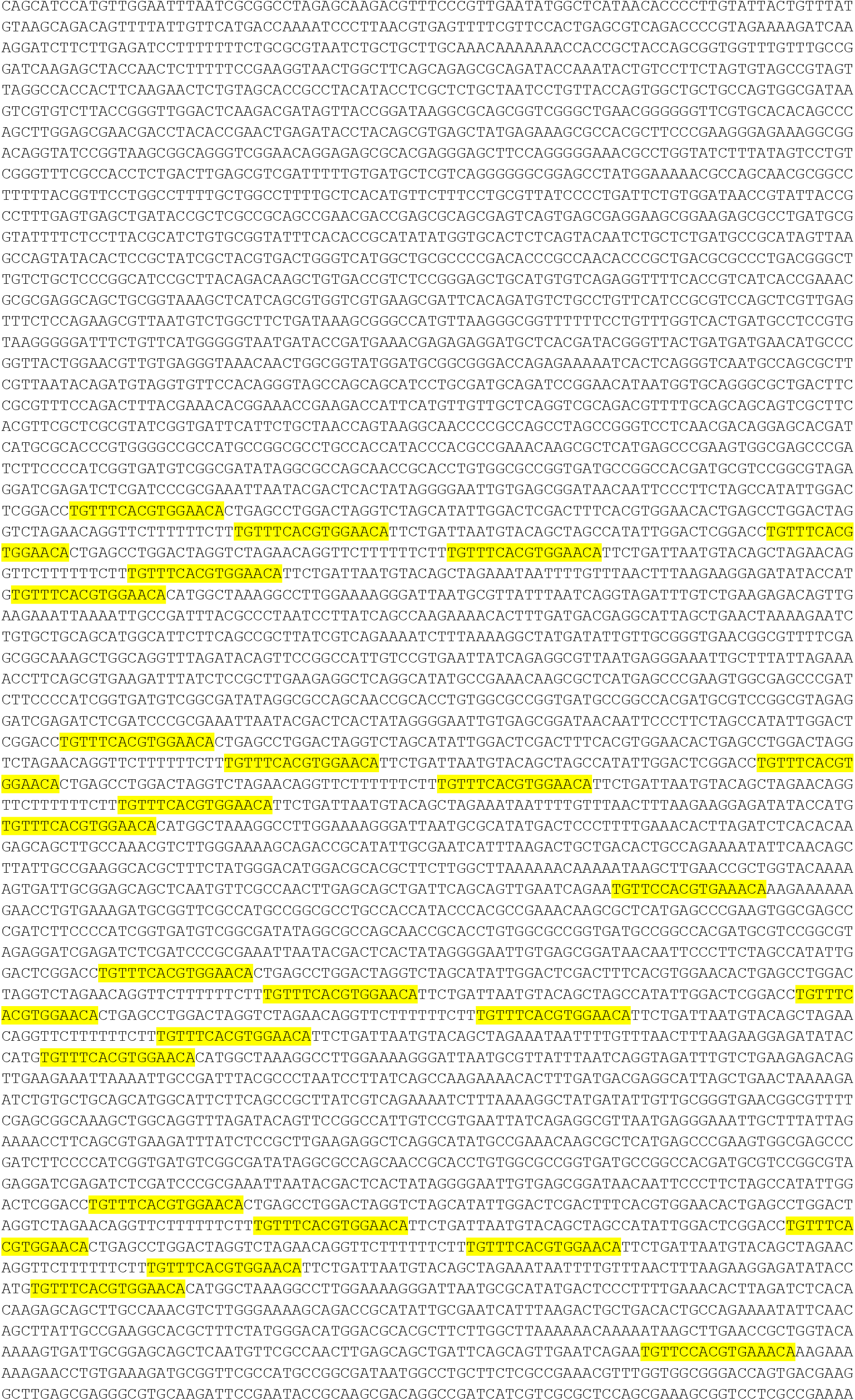
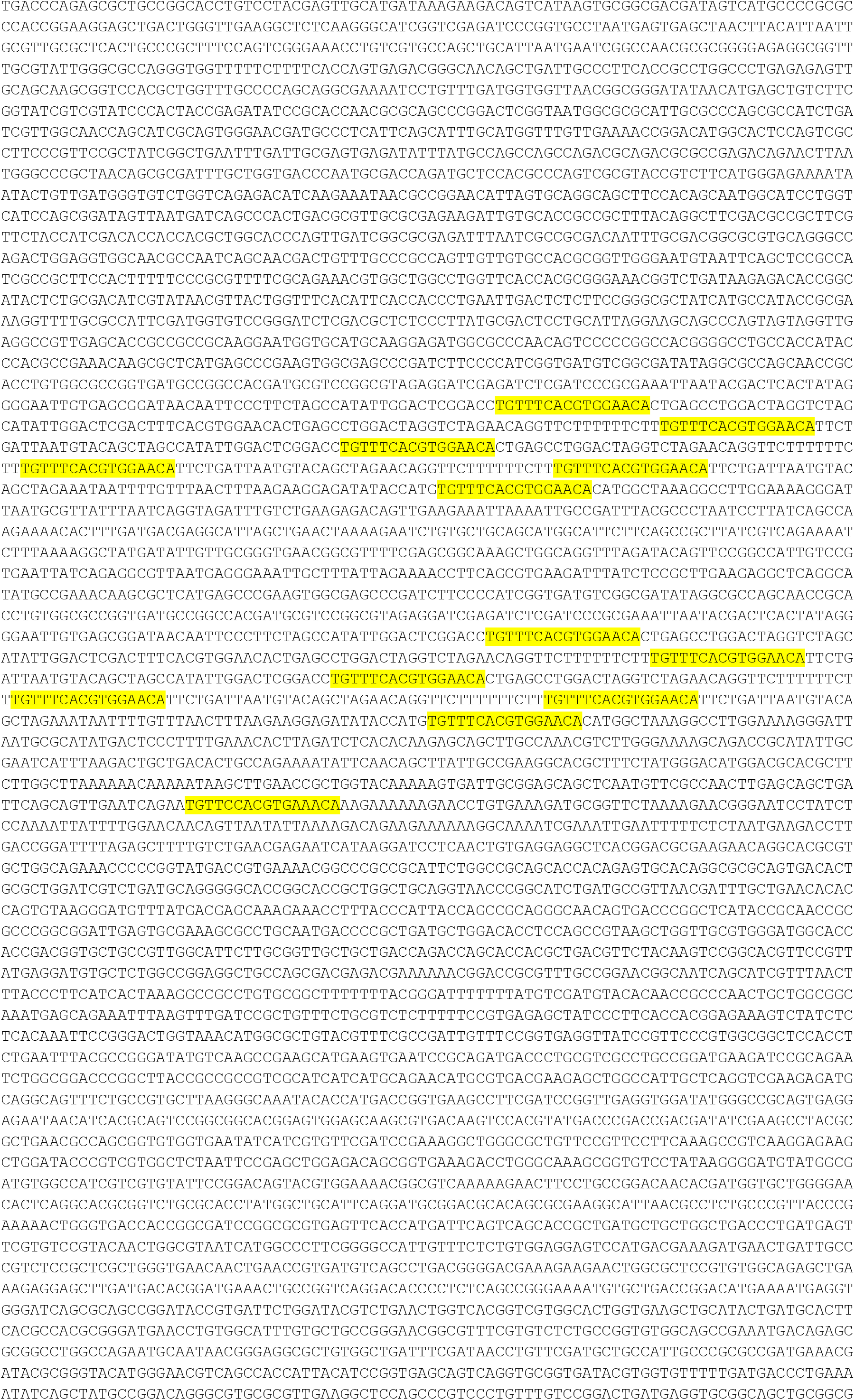
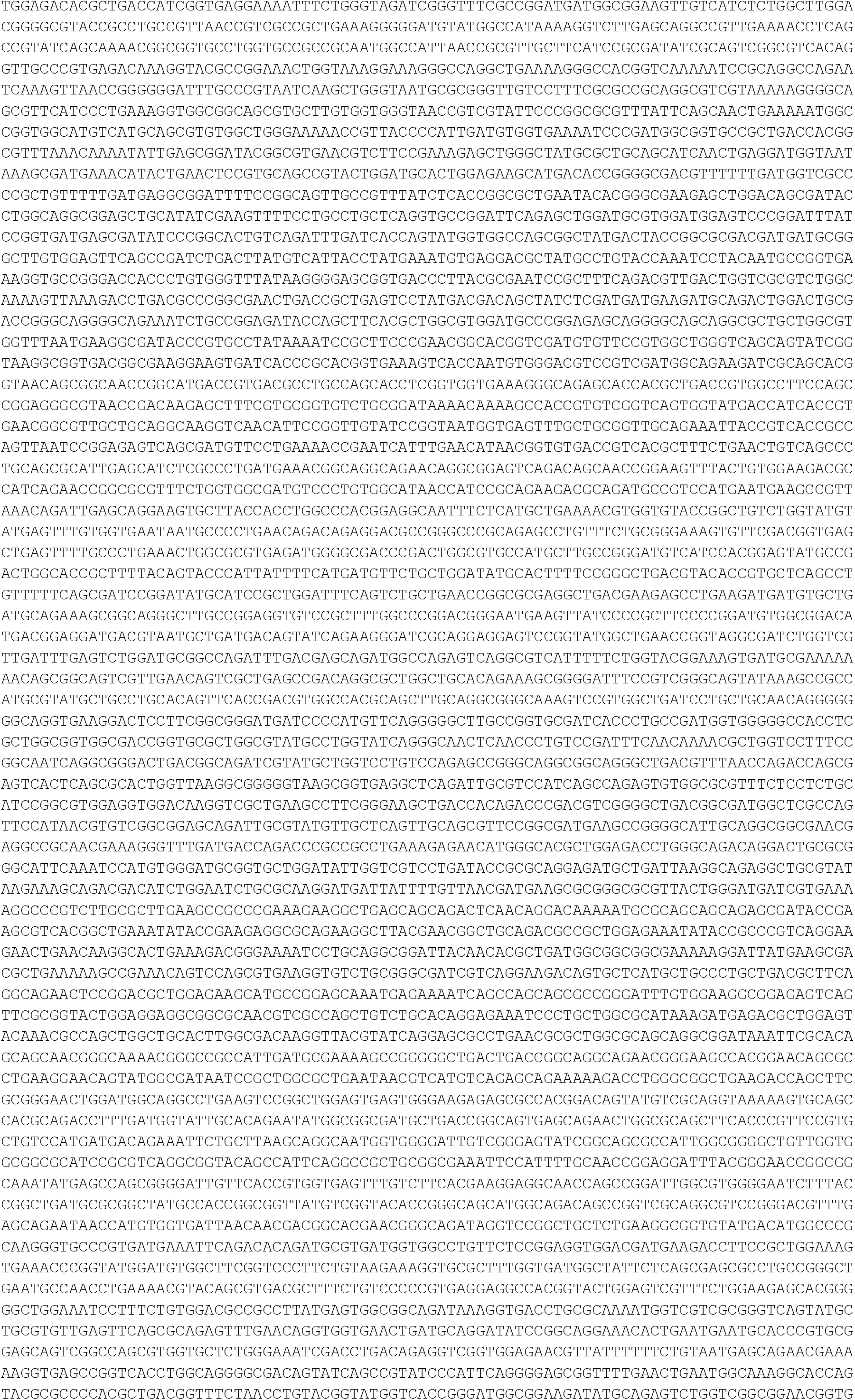
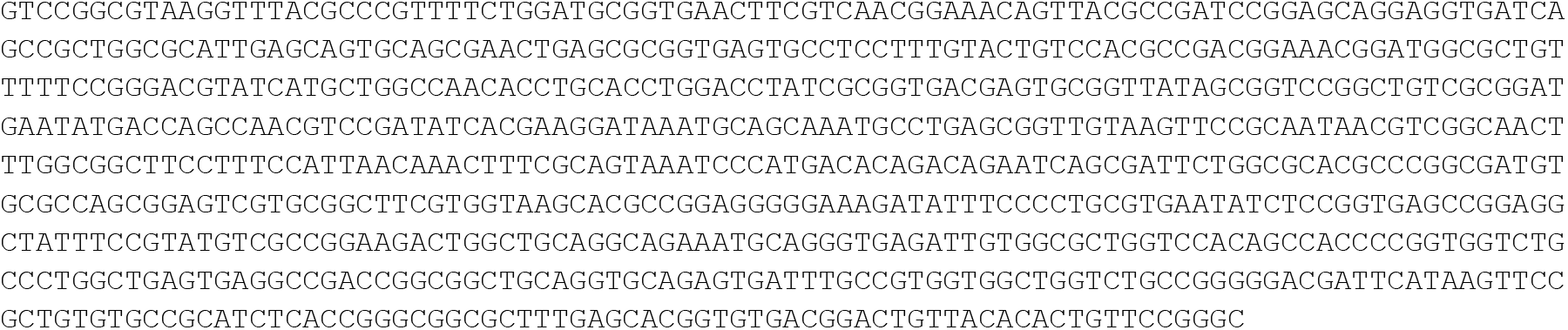

#### C-trap EcoRI 39x parS DNA (17344 bp)

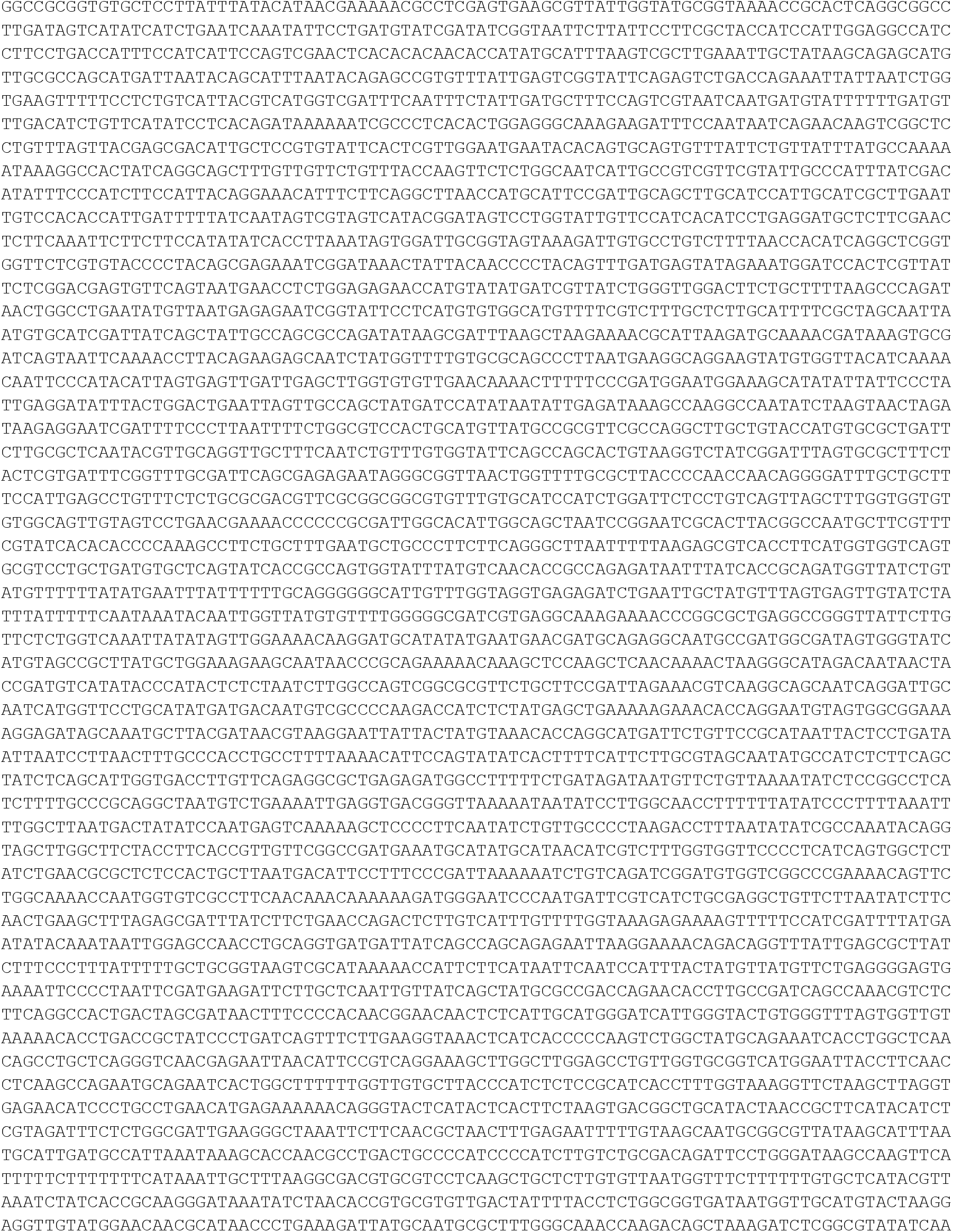
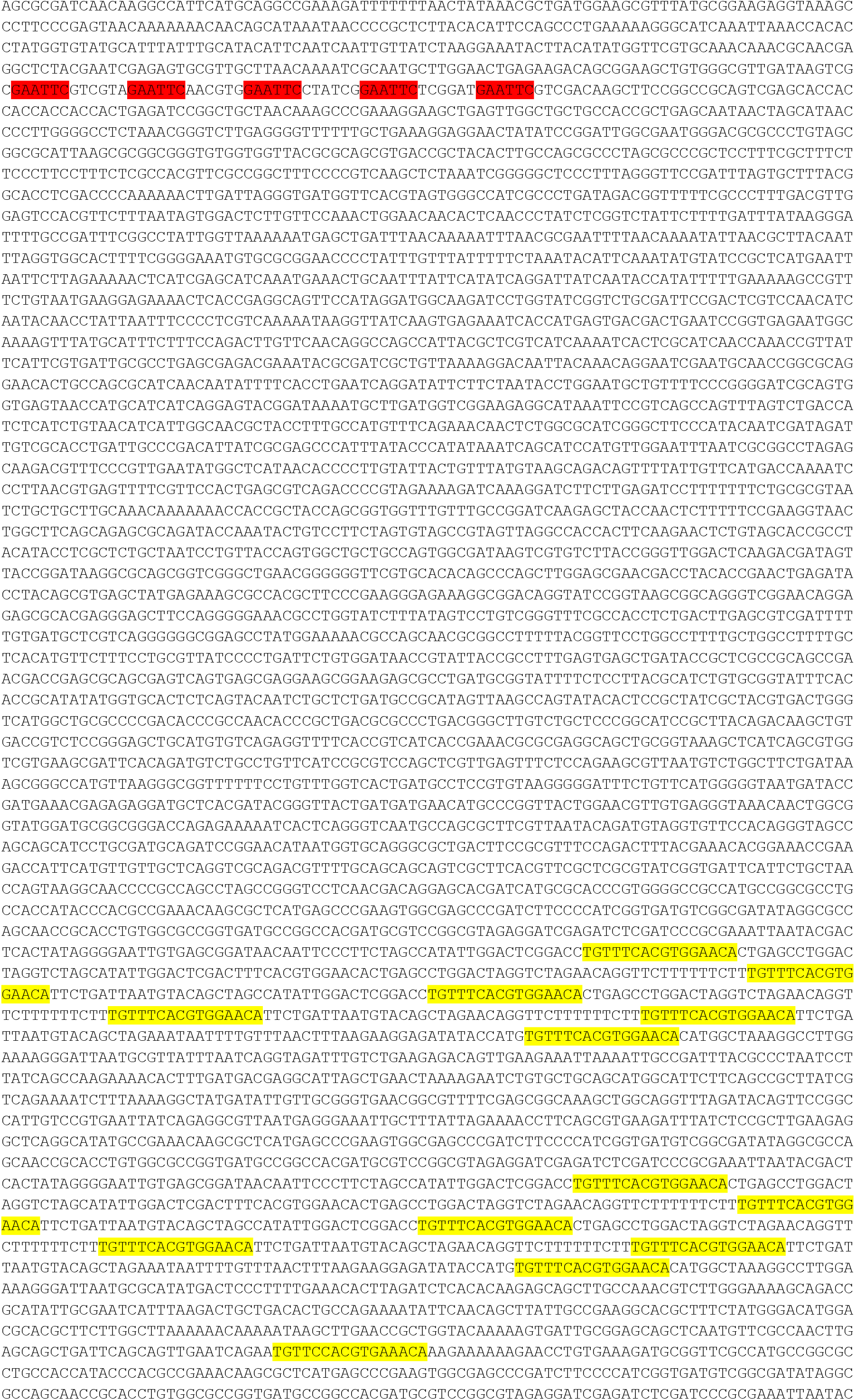
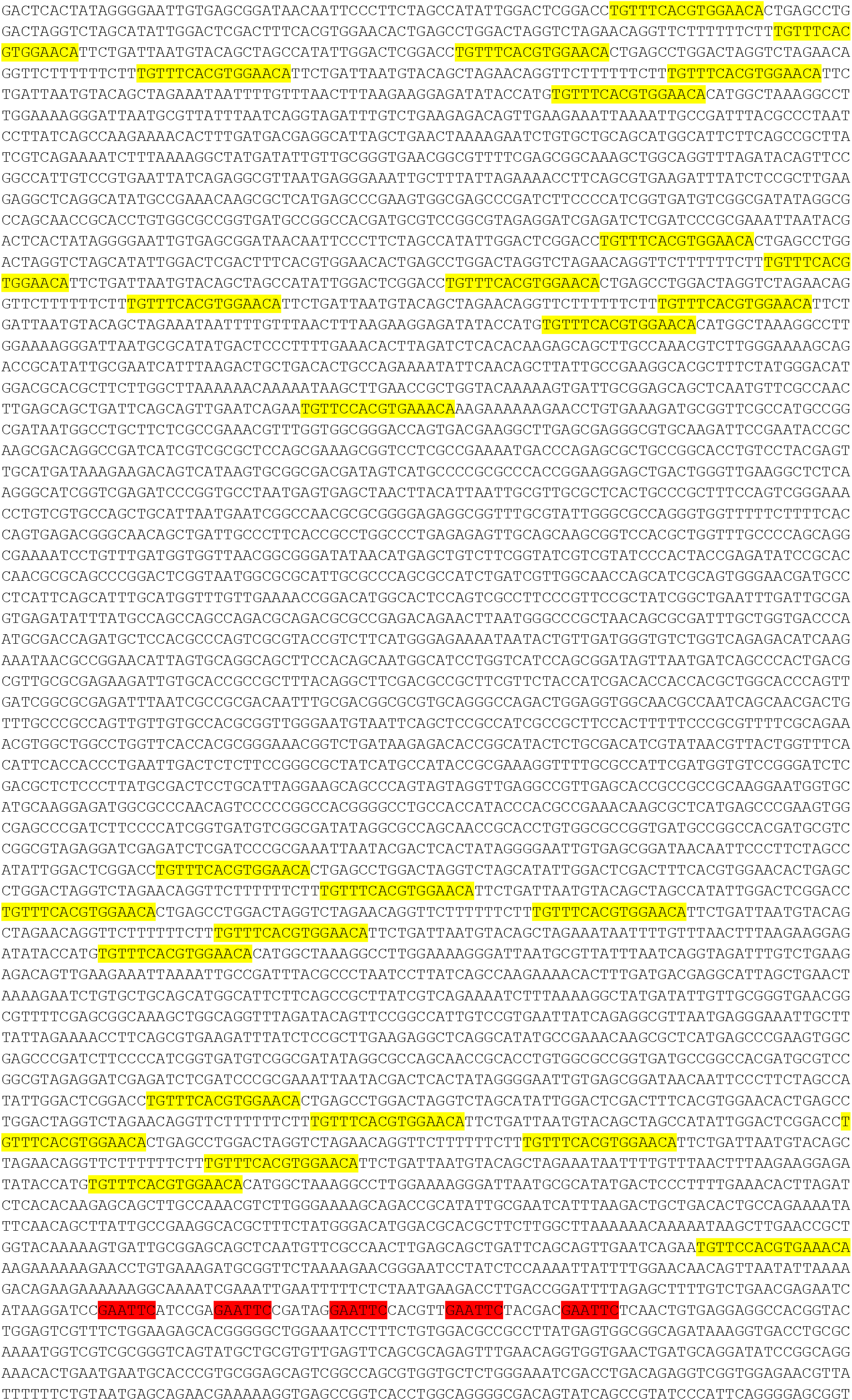
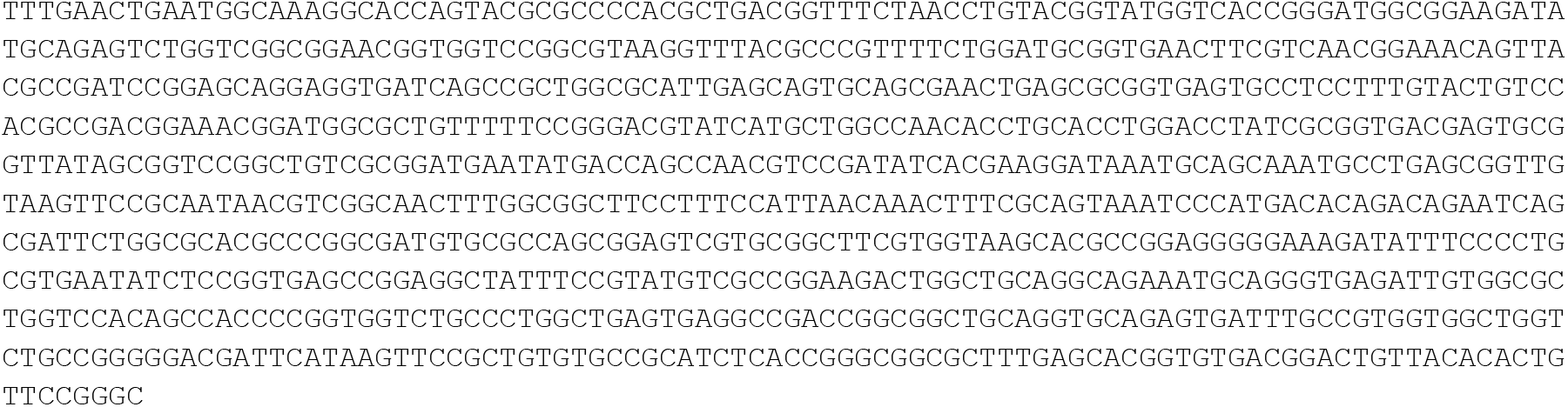

## REFERENCES

Autret S, Nair R, Errington J. 2001. Genetic analysis of the chromosome segregation protein Spo0J of Bacillus subtilis: evidence for separate domains involved in DNA binding and interactions with Soj protein. Mol Microbiol 41:743–755.

Bartosik AA, Lasocki K, Mierzejewska J, Thomas CM, Jagura-Burdzy G. 2004. ParB of Pseudomonas aeruginosa: interactions with its partner ParA and its target parS and specific effects on bacterial growth. J Bacteriol 186:6983–6998. doi:10.1128/JB.186.20.6983-6998.2004

Bouet JY, Funnell BE. 1999. P1 ParA interacts with the P1 partition complex at parS and an ATP-ADP switch controls ParA activities. EMBO J 18:1415–1424. doi:10.1093/emboj/18.5.1415

Breier AM, Grossman AD. 2007. Whole-genome analysis of the chromosome partitioning and sporulation protein Spo0J (ParB) reveals spreading and origin-distal sites on the Bacillus subtilis chromosome. Mol Microbiol 64:703–718. doi:10.1111/j.1365-2958.2007.05690.x

Broedersz CP, Wang X, Meir Y, Loparo JJ, Rudner DZ, Wingreen NS. 2014. Condensation and localization of the partitioning protein ParB on the bacterial chromosome. Proc Natl Acad Sci U S A 111:8809–8814. doi:10.1073/pnas.1402529111

Brüning J-G, Howard JAL, Myka KK, Dillingham MS, McGlynn P. 2018. The 2B subdomain of Rep helicase links translocation along DNA with protein displacement. Nucleic Acids Res 46:8917–8925. doi:10.1093/nar/gky673

Candelli A, Wuite GJ, Peterman EJ. 2011. Combining optical trapping, fluorescence microscopy and micro-fluidics for single molecule studies of DNA-protein interactions. Phys Chem Chem Phys 13:7263–7272. doi:10.1039/c0cp02844d

Carrasco C, Gilhooly NS, Dillingham MS, Moreno-Herrero F. 2013. On the mechanism of recombination hotspot scanning during double-stranded DNA break resection. Proc Natl Acad Sci U S A 110:E2562–E2571. doi:10.1073/pnas.1303035110

Chen BW, Lin MH, Chu CH, Hsu CE, Sun YJ. 2015. Insights into ParB spreading from the complex structure of Spo0J and parS. Proc Natl Acad Sci U S A 112:6613–6618. doi:10.1073/pnas.1421927112

Davis MA, Martin KA, Austin SJ. 1992. Biochemical activities of the ParA partition protein of the P1 plasmid. Mol Microbiol 6:1141–1147. doi:10.1111/j.1365-2958.1992.tb01552.x

Fisher GLM, Pastrana CL, Higman VA, Koh A, Taylor JA, Butterer A, Craggs T, Sobott F, Murray H, Crump MP, Moreno-Herrero F, Dillingham MS. 2017. The structural basis for dynamic DNA binding and bridging interactions which condense the bacterial centromere. Elife 6:e28086. doi:10.7554/eLife.28086

Funnell BE. 2016. ParB Partition Proteins: Complex Formation and Spreading at Bacterial and Plasmid Centromeres. Front Mol Biosci 3:1–6. doi:10.3389/fmolb.2016.00044

Graham TG, Wang X, Song D, Etson CM, van Oijen AM, Rudner DZ, Loparo JJ. 2014. ParB spreading requires DNA bridging. Genes Dev 28:1228–1238. doi:10.1101/gad.242206.114

Gruber S, Errington J. 2009. Recruitment of condensin to replication origin regions by ParB/SpoOJ promotes chromosome segregation in B. subtilis. Cell 137:685–696. doi:10.1016/j.cell.2009.02.035

Guilhas B, Walter J-C, Rech J, David G, Walliser NO, Palmeri J, Mathieu-Demaziere C, Parmeggiani A, Bouet J-Y, Le Gall A, Nollmann M. 2020. ATP-Driven Separation of Liquid Phase Condensates in Bacteria. Mol Cell 79:293-303.e4. doi:https://doi.org/10.1016/j.molcel.2020.06.034

Gutierrez-Escribano P, Hormeño S, Madariaga-Marcos J, Sole-Soler R, O’Reilly FJ, Morris K, Aicart-Ramos C, Aramayo R, Montoya A, Kramer H, Rappsilber J, Torres-Rosell J, Moreno-Herrero F, Aragon L. 2020. Purified Smc5/6 complex exhibits DNA substrate recognition and compaction. Mol Cell (in press).

Hayes F, Barillà D. 2006. The bacterial segrosome: a dynamic nucleoprotein machine for DNA trafficking and segregation. Nat Rev Microbiol 4:133–143. doi:10.1038/nrmicro1342

Horcas I, Fernández R, Gómez-Rodríguez JM, Colchero J, Gómez-Herrero J, Baro AM. 2007. WSXM: a software for scanning probe microscopy and a tool for nanotechnology. Rev Sci Instrum 78:013705. doi:10.1063/1.2432410

Jalal AS, Tran NT, L. TB. 2020. ParB spreading on DNA requires cytidine triphosphate in vitro. Elife 9:e53515. doi:10.7554/eLife.53515

Jecz P, Bartosik AA, Glabski K, Jagura-Burdzy G. 2015. A single parS sequence from the cluster of four sites closest to oriC is necessary and sufficient for proper chromosome segregation in Pseudomonas aeruginosa. PLoS One. doi:10.1371/journal.pone.0120867

Kemmerich FE, Kasaciunaite K, Seidel R. 2016. Modular magnetic tweezers for single-molecule characterizations of helicases. Methods 108:4–13. doi:10.1016/j.ymeth.2016.07.004

King K, Benkovic SJ, Modrich P. 1989. Glu-111 is required for activation of the DNA cleavage center of EcoRI endonuclease. J Biol Chem 264:11807–11815.

Kusiak M, Gapczynska A, Plochocka D, Thomas CM, Jagura-Burdzy G. 2011. Binding and spreading of ParB on DNA determine its biological function in Pseudomonas aeruginosa. J Bacteriol 193:3342–3355. doi:10.1128/JB.00328-11

Leonard TA, Butler PJ, Lowe J. 2004. Structural analysis of the chromosome segregation protein Spo0J from Thermus thermophilus. Mol Microbiol 53:419–432. doi:10.1111/j.1365-2958.2004.04133.x

Lin DC, Grossman AD. 1998. Identification and characterization of a bacterial chromosome partitioning site. Cell 92:675–685.

Madariaga-Marcos J, Hormeño S, Pastrana CL, Fisher GLM, Dillingham MS, Moreno-Herrero F. 2018. Force determination in lateral magnetic tweezers combined with TIRF microscopy. Nanoscale 10:4579–4590. doi:10.1039/c7nr07344e

Madariaga-Marcos J, Pastrana CL, Fisher GLM, Dillingham MS, Moreno-Herrero F. 2019. ParB dynamics and the critical role of the CTD in DNA condensation unveiled by combined force-fluorescence measurements. Elife 8:e43812. doi:10.7554/eLife.43812

Marbouty M, Le Gall A, Cattoni DI, Cournac A, Koh A, Fiche JB, Mozziconacci J, Murray H, Koszul R, Nollmann M. 2015. Condensin- and Replication-Mediated Bacterial Chromosome Folding and Origin Condensation Revealed by Hi-C and Super-resolution Imaging. Mol Cell 59:588–602. doi:10.1016/j.molcel.2015.07.020

Minnen A, Bürmann F, Wilhelm L, Anchimiuk A, Diebold-Durand ML, Gruber S. 2016. Control of Smc Coiled Coil Architecture by the ATPase Heads Facilitates Targeting to Chromosomal ParB/parS and Release onto Flanking DNA. Cell Rep 14:2003–2016. doi:10.1016/j.celrep.2016.01.066

Murray H, Ferreira H, Errington J. 2006. The bacterial chromosome segregation protein Spo0J spreads along DNA from parS nucleation sites. Mol Microbiol 61:1352–1361. doi:10.1111/j.1365-2958.2006.05316.x

Newton MD, Taylor BJ, Driessen RPC, Roos L, Cvetesic N, Allyjaun S, Lenhard B, Cuomo ME, Rueda DS. 2019. DNA stretching induces Cas9 off-target activity. Nat Struct Mol Biol 26:185–192. doi:10.1038/s41594-019-0188-z

Osorio-Valeriano M, Altegoer F, Steinchen W, Urban S, Liu Y, Bange G, Thanbichler M. 2019. ParB-type DNA Segregation Proteins Are CTP-Dependent Molecular Switches. Cell 179:1512–1524. doi:10.1016/j.cell.2019.11.015

Pastrana CL, Carrasco C, Akhtar P, Leuba SH, Khan SA, Moreno-Herrero F. 2016. Force and twist dependence of RepC nicking activity on torsionally-constrained DNA molecules. Nucleic Acids Res 44:8885–8896. doi:10.1093/nar/gkw689

Radnedge L, Youngren B, Davis M, Austin S. 1998. Probing the structure of complex macromolecular interactions by homolog specificity scanning: The P1 and P7 plasmid partition systems. EMBO J 17:6076–85. doi:10.1093/emboj/17.20.6076

Rodionov O, ŁObocka M, Yarmolinsky M. 1999. Silencing of genes flanking the P1 plasmid centromere. Science 283:546–549. doi:10.1126/science.283.5401.546

Rodionov O, Yarmolinsky M. 2004. Plasmid partitioning and the spreading of P1 partition protein ParB. Mol Microbiol 52:1215–1223. doi:10.1111/j.1365-2958.2004.04055.x

Sanchez A, Cattoni DI, Walter JC, Rech J, Parmeggiani A, Nollmann M, Bouet JY. 2015. Stochastic Self-Assembly of ParB Proteins Builds the Bacterial DNA Segregation Apparatus. Cell Syst 1:163–173. doi:10.1016/j.cels.2015.07.013

Schumacher MA. 2008. Structural biology of plasmid partition: uncovering the molecular mechanisms of DNA segregation. Biochem J 412:1–18. doi:10.1042/BJ20080359

Schumacher MA, Funnell BE. 2005. Structures of ParB bound to DNA reveal mechanism of partition complex formation. Nature 438:516–519. doi:10.1038/nature04149

Soh YM, Davidson IF, Zamuner S, Basquin J, Bock FP, Taschner M, Veening JW, de Los Rios P, Peters JM, Gruber S. 2019. Self-organization of parS centromeres by the ParB CTP hydrolase. Science 366:1129–1133. doi:10.1126/science.aay3965

Song D, Loparo JJ. 2015. Building bridges within the bacterial chromosome. Trends Genet 31:164–173. doi:10.1016/j.tig.2015.01.003

Song D, Rodrigues K, Graham TGW, Loparo JJ. 2017. A network of cis and trans interactions is required for ParB spreading. Nucleic Acids Res 45:7106–7117. doi:10.1093/nar/gkx271

Strick TR, Allemand JF, Bensimon D, Croquette V. 1998. Behavior of supercoiled DNA. Biophys J 74:2016–28. doi:10.1016/S0006-3495(98)77908-1

Sullivan NL, Marquis KA, Rudner DZ. 2009. Recruitment of SMC by ParB-parS organizes the origin region and promotes efficient chromosome segregation. Cell 137:697–707. doi:10.1016/j.cell.2009.04.044

Taylor JA, Pastrana CL, Butterer A, Pernstich C, Gwynn EJ, Sobott F, Moreno-Herrero F, Dillingham MS. 2015. Specific and non-specific interactions of ParB with DNA: Implications for chromosome segregation. Nucleic Acids Res 43:719–731. doi:10.1093/nar/gku1295

Vecchiarelli AG, Han Y-W, Tan X, Mizuuchi M, Ghirlando R, Biertümpfel C, Funnell BE, Mizuuchi K. 2010. ATP control of dynamic P1 ParA-DNA interactions: a key role for the nucleoid in plasmid partition. Mol Microbiol 78:78–91. doi:10.1111/j.1365-2958.2010.07314.x

Walter J-C, Rech J, Walliser N-O, Dorignac J, Geniet F, Palmeri J, Parmeggiani A, Bouet J-Y. 2020. Physical Modeling of a Sliding Clamp Mechanism for the Spreading of ParB at Short Genomic Distance from Bacterial Centromere Sites. iScience 23:101861. doi:https://doi.org/10.1016/j.isci.2020.101861

Wang X, Brandão HB, Le TBK, Laub MT, Rudner DZ. 2017. Bacillus subtilis SMC complexes juxtapose chromosome arms as they travel from origin to terminus. Science 355:524–527. doi:10.1126/science.aai8982

Wang X, Tang OW, Riley EP, Rudner DZ. 2014. The SMC condensin complex is required for origin segregation in Bacillus subtilis. Curr Biol 24:287–292. doi:10.1016/j.cub.2013.11.050

Wasserman MR, Schauer GD, O’Donnell ME, Liu S. 2020. Replication Fork Activation is Enabled by a Single-Stranded DNA Gate in CMG Helicase. Biophys J. doi:10.1016/j.bpj.2019.11.2143

